# Matrix stiffness controls ciliogenesis and centriole position

**DOI:** 10.1101/2021.08.12.452439

**Authors:** Ivanna Williantarra, Sophia Leung, Yu Suk Choi, Ashika Channa, Sue R McGlashan

**Affiliations:** Department of Anatomy and Medical Imaging, Faculty of Medical and Health Sciences, University of Auckland, New Zealand; School of Human Sciences, University of Western Australia, Australia

**Keywords:** primary cilium, mechanobiology, substrate stiffness, temporal dynamics, actin, centriole positioning

## Abstract

Mechanical stress and the stiffness of the extracellular matrix are key drivers of tissue development and homeostasis. Aberrant mechanosensation is associated with a wide range of pathologies, including diseases such as osteoarthritis. Substrate stiffness is one of the well-known mechanical properties of the matrix that enabled establishing the central dogma of an integrin-mediated mechanotransduction using stem cells. However, how specific cells ‘feel’ or sense substrate stiffness requires further study. The primary cilium is an essential cellular organelle that senses and integrates mechanical and chemical signals from the extracellular environment. We hypothesised that the primary cilium dynamically alters its length and position to fine-tune cell mechanosignalling based on substrate stiffness alone. We used a hydrogel system of varying substrate stiffness to examine the role of substrate stiffness on cilia frequency, length and centriole position as well as cell and nuclei area over time. Contrary to other cell types, we show that chondrocyte primary cilia shorten on softer substrates demonstrating tissue-specific mechanosensing which is aligned with the tissue stiffness the cells originate from. We further show that stiffness alone determines centriole positioning to either the basal or apical membrane during attachment and spreading, with centriole positioned towards the basal membrane on stiffer substrates. These phenomena are mediated by force generation actin-myosin stress fibres in a time-dependent manner. Based on these findings, we propose that substrate stiffness plays a central role in cilia positioning, regulating cellular response to external forces, and may be a key driver of mechanosignalling-associated diseases.

**Significance Statement:** The primary cilium has been thrust into the limelight owing to its role as a cellular sensor in embryonic development and adult tissue maintenance. How the primary cilium interacts with the mechanical environment still remains unclear. We show that substrate stiffness dynamically regulates primary cilium length and position through integrin-mediated traction forces, the cilia are a key determinant of cell shape on certain stiffnesses. Our data support the promising potential of primary cilia as a novel target in mechanotherapy for improved clinical outcomes in cartilage pathobiology.

## Introduction

Matrix (or substrate) stiffness is a major factor in cell spreading [1], [2], adhesion [3], proliferation and differentiation [4]. Mimicking the stiffness of neural, muscle and bone tissues, substrate stiffness induced differentiation toward those specific tissues [4], [5]. In articular cartilage, a specialised tissue present in synovial joints, the stiff extracellular matrix (ECM) distributes stresses and reduces friction between joints. Chondrocytes, the only cell type present in cartilage, are solely responsible for the production and maintenance of the ECM and in part, the local mechanical tissue microenvironment regulates this process. In joint diseases such as osteoarthritis (OA), the ECM becomes softer, loses its protective abilities, leading to altered chondrocyte metabolism, inflammation and cartilage degeneration [6]–[8]. The precise mechanisms of how chondrocytes sense the environment and how this may go awry in OA is not well understood. Therefore, this study examines how the stiffness of the extracellular microenvironment influences cellular mechanosensitivity through the action of the primary cilium.

The primary cilium is a microtubule-based signalling organelle present on almost all cell types, including chondrocytes. They act as cellular sensors, receiving and integrating diverse signals such as morphogens, growth factors and mechanical stimuli from the extracellular environment in a tissue-specific manner [9]. Unlike primary cilia of epithelial cells that protrude in a lumen, chondrocyte cilia are closely associated with the tensile collagen fibres in cartilage. They express collagen-binding integrins and are mechanically deflected through ECM interactions [10]–[13]. Chondrocyte cilia are enriched with mechanoreceptors such as integrins, G protein-couple receptors, and calcium channels [39], [40]. Compressive or tensile forces modulate cilia frequency and length, and removal of cilia results in reduced mechanotransduction [14]–[17]. In the absence of primary cilia, articular cartilage’s structure and mechanical properties are dramatically altered [18].

The process of cilia assembly is termed ciliogenesis. Cells assemble a cilium following cell division [19]. Upon assembly, cilia adjust in length in response to chemical and mechanical stimuli, and in turn, can enhance signal detection [20], [21]. Conversely, cilia shortening or resorption decreases responsiveness to signals, providing a mechanism of sensory signal adaptation [14], [22]. The cilia position within tissues also plays a key role in development and homeostasis [23], [24]. Polarised cells place primary cilia on the apical surface, commonly protruding into a lumen, e.g. kidney epithelial cell cilia sense flow and metabolites within urine [21], [25]. In contrast, cilia show tissue-specific position and length in 3D culture of non-polarised cells, such as chondrocytes. Cilia are oriented in specific planes in both articular cartilage and the epiphyseal growth plate, where they coordinate proliferation and growth in the developing long bone [26]–[28]. Chondrocyte primary cilia length also varies with cartilage depth and with osteoarthritis [29].

The role of the microenvironment in cellular mechanosensation and tissue-specific behaviour is commonly studied *in vitro* using engineered matrices of tunable stiffness [4]. It is well established that cells cultured on stiff substrates with Young’s modulus >25 kPa show increased cell spreading and adhesion with a significant stress fibre network comprised of filamentous actin (F-actin) [5], [30]. Stress fibers generate traction forces and actomyosin network contraction in increased ECM stiffness [3], [31]–[34]. Recent studies have revealed that the actin cytoskeleton and regulators of actin dynamics also modulate cilia assembly. Actin perturbation using cytochalasin D [35] increases ciliogenesis and cilia length. However, limited studies have deeply probed the relationships between primary cilia and the better known actin cytoskeleton in chondrocytes. We believe that the primary cilium is the missing link connecting and integrating these phenomena.

In this study, we use a tuneable polyacrylamide gel system with stiffness representing healthy and degenerated cartilage ECM to understand the role of substrate stiffness in ciliogenesis and cilia length over time. We show an in-depth understanding of how the mechanical environment alone regulates primary cilia formation and position and potentially enhancing or limiting access to key signalling ligands and/or mechanical stimuli.

## Results

### Stiff substrates increase cilia length but not frequency

To investigate the effect of substrate stiffness on primary cilia, we examined cilia frequency and length of cells cultured on polyacrylamide gel substrates with stiffnesses of 5 kPa (4.92 ± 0.08 kPa) and 50 kPa (50.10 ± 0.89 kPa), as well as glass coverslips (>1 GPa). Primary cilia and centrioles were detected using antibodies against ARL13B and γ-tubulin in cells cultured on all three substrates (figure 1A). Media containing 10% serum was removed and replaced with 0.25% serum (time = 0 h) for 48h and hereinafter is referred to as the serum starvation period. The percentage of ciliated cells 12 hours prior to the starvation period (pre-starvation in figure 1B) was similar on all stiffness (22%, 24% and 26% on 5 kPa, 50 kPa and coverslips, respectively). As expected, serum starvation for 48 h resulted in greater cilia frequency on all three substrates, but stiffness did not affect cilia frequency at any time point (p>0.05). Stiffness did, however, significantly modulate cilia length. Cilia length on stiffer substrates was greatest at 12 h (p<0.001, figure 1C) and this trend was maintained for the remaining culture period. Cilia of cells cultured on coverslips were approximately 0.5 μm longer than cilia of cells on 5 kPa gels at all time points (p=0.035 at 12 h, p<0.001 at 24 h and 48 h). We next examined the effect of each stiffness on cilia length during serum starvation. Interestingly, on the two extreme stiffnesses (5 kPa and coverslips), cilia length remained unchanged during serum starvation (p>0.05). However, on 50 kPa gels, cilia showed transient changes in cilia length and were longest at 24 h following serum removal (p<0.001).

**Figure 1.**
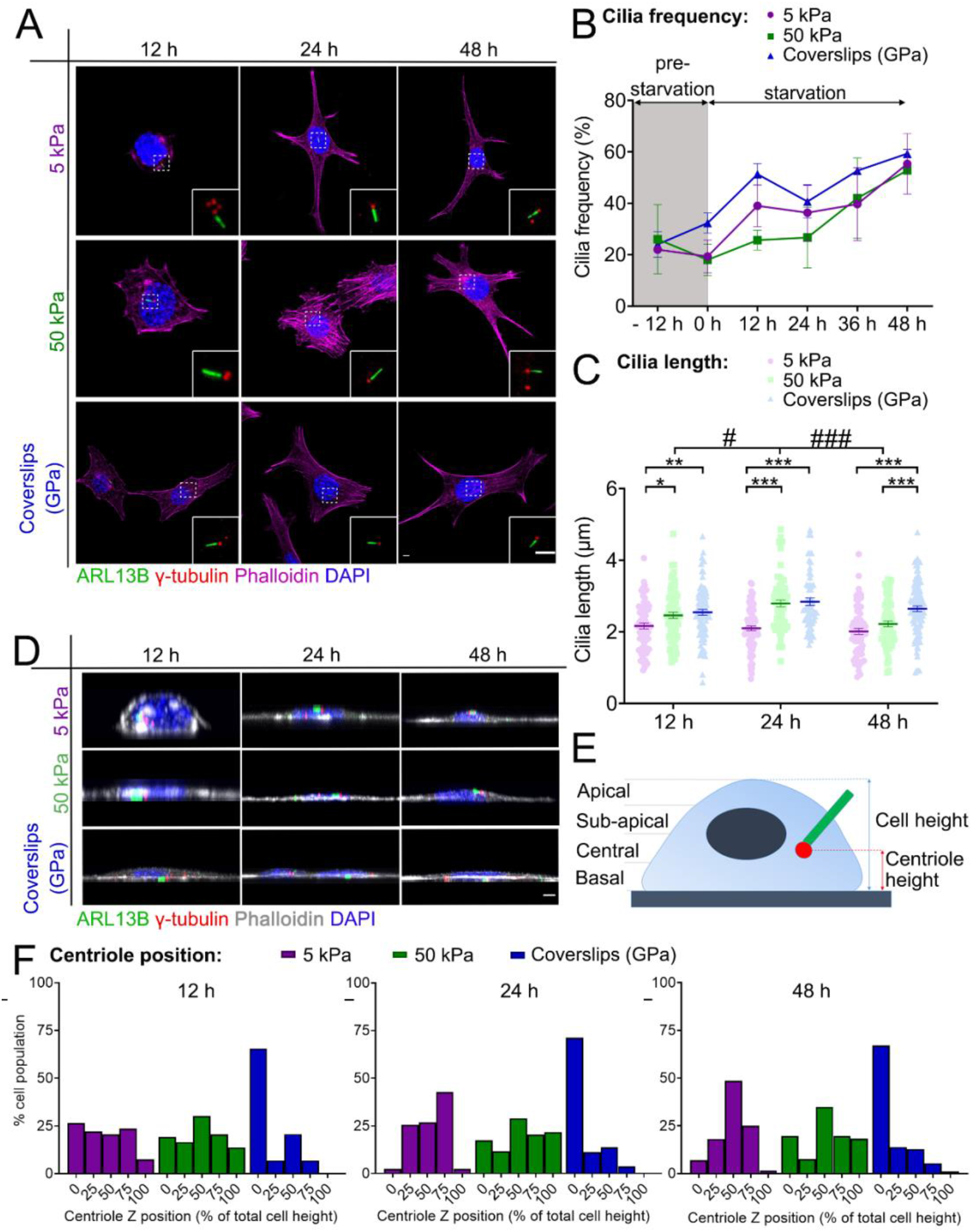
The effect of substrate stiffness on primary cilia characteristics (A) Confocal images of ciliated chondrocytes cultured on 5 kPa, 50 kPa and coverslips (GPa) at different time points following serum starvation. Immunostaining for the primary cilia (ARL13B: green), centriole (γ-tubulin: red), actin (phalloidin: magenta) and nucleus (DAPI: blue). White dashed boxes show the presence of primary cilia. Insets show a magnified view of primary cilia and centriole. Scale bar = 5 μm. Transient changes in (B) cilia frequency and (C) cilia length as a function of stiffness and time. (D) XZ projections of ciliated chondrocytes cultured on at 12h, 24h and 48h after starvation. Primary cilia (ARL13B: green), centriole (γ-tubulin: red), actin (phalloidin: grey) and nucleus (DAPI: blue). Scale bar = 5 μm. (E) Schematic Z stack images of centriole positioning and cell height (F) The frequency distribution histogram of centriole position (as a percentage of cell height) for cells cultured on 5 kPa, 50 kPa hydrogels and glass coverslips following serum starvation at 12 h, 24 h and 48 h. 0 h refers to the start of the starvation period. Values represent mean ± SEM. For cilia frequency, N = 3; n ≥100 cilia per stiffness. A total number of 900 cells were examined. For cilia length and centriole position, N=3; n≥66. A total number of 690 cells were examined. * for comparison between stiffness and # for comparison between time points. *p<0.05, **p<0.01, ***p<0.001.

### Substrate stiffness determines centriole positioning

We next examined the effect of stiffness on centriole positioning and cilia orientation. For the first time, we showed that stiffness alone can direct centrioles to particular regions within a cell and that centrioles are more likely to be located towards the basal surface on stiffer substrates. Figure 1D shows representative side-on view images of centriole position relative to cell height on different stiffnesses over time. At 12 h, in cells cultured on 5 and 50 kPa, centrioles were distributed relatively evenly across the XZ cell view (figure 1F). However, at 24 h on 5 kPa gels, 43% of centrioles were located within the subapical domain and only 3% at the apical domain (p=0.012 compared to 12 h). In contrast, cells on 50 kPa gels showed an even centriole distribution at all time points (p>0.05) and centrioles in cells cultured on coverslips were positioned close to the basal surface at all time points (p>0.05). We also investigated the effect of substrate stiffness on cilia orientation by measuring cilia elevation angle (figure S1A). Overall, the distribution of the elevation angle peaked near 0°, indicating that cilia were primarily within the XY plane. Further, cilia orientation was not significantly influenced by substrate stiffness or duration of serum starvation (p=0.129 for the effect of stiffness and p=0.422 for the effect of time) (figure S1B).

### Substrate stiffness influences cell morphology and actin architecture

It is well known that substrate stiffness strongly influences cell morphology [36], [37] so we next examined the cell morphology of ciliated cells cultured on different stiffnesses. In general, independent of time, cells on 5 kPa gels showed less organised stress fibre distribution, with actin puncta distributed across the whole cell volume (figure 2A). In contrast, with increasing stiffness, an associated increase in cell area (p<0.001) and F-actin fibres was more prominent, as shown in Figures 2B and 2G. Each stiffness yielded varied temporally dynamic and stiffness-dependent changes in the cell area. Cell area on 5 kPa gels reduced over time (p<0.001 at 24 h and 48 h) while cells on coverslips showed a transient increase at 24h (p=0.001), which then reduced at 48 h (p<0.001). In contrast, cell area on 50 kPa stiffness gels had no effect on cell area over time (p>0.05). Figure 2C shows that stiffer substrates were associated with an increase in the nucleus area (p<0.001) on both the soft and coverslips but again, no major effect of 50 kPa was observed (p>0.05).

**Figure 2.**
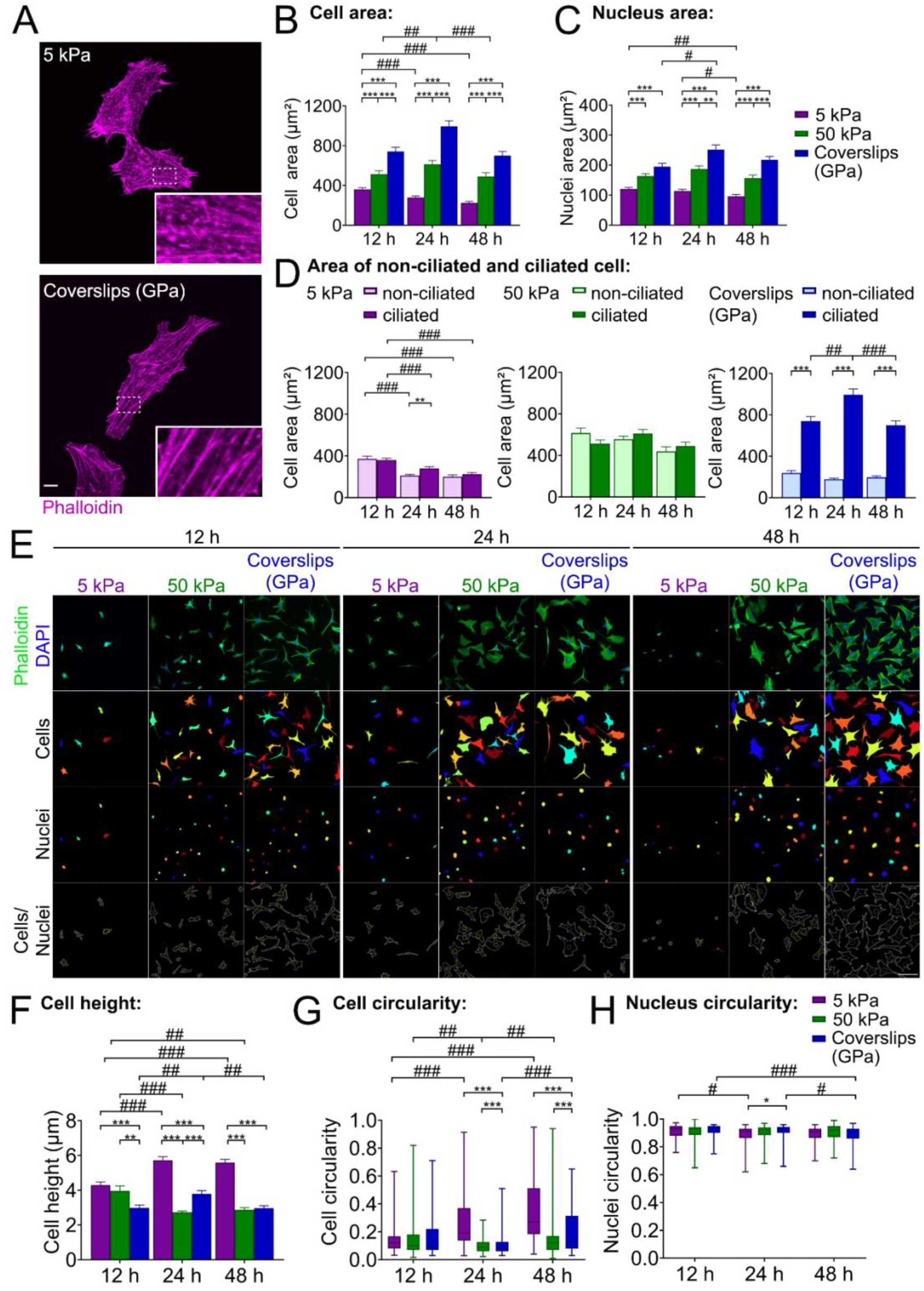
(A) Representative confocal images of mouse articular chondrocytes stained for F-actin with phalloidin on 5 kPa gels and coverslips. Scale bar = 10 μm. (B) Cell area and (C) nucleus area as a function of substrate stiffness over time in ciliated cells. (D) Difference of cell area between non-ciliated and ciliated cells cultured on 5 kPa, 50 kPa and coverslips. (E) Representative images showing differences in size and circularity of chondrocytes cultured on 5 kPa, 50 kPa and coverslips at different time points. Images were taken at 20? magnification using phalloidin for cytoplasmic and DAPI for nuclear boundary recognition. Scale bar = 100 μm. Cell morphology as a function of substrate stiffness over time: (F) cell height; (G) cell circularity and (H) nucleus circularity. For ciliated cells, N=3; n≥66 per condition. A total number of 690 cells were examined. For non-ciliated cells, N=3; n ≥62 cells per condition. A total number of 689 cells were examined. * for comparison between stiffness and # for comparison between time points.*p<0.05, **p<0.01, ***p<0.001.

To determine if these cell morphological phenomena were cilia-dependent, we next examined how substrate stiffness influenced cell area in non-ciliated cells (figure 2D). Strikingly, non-ciliated cells cultured on coverslips had significantly smaller cell areas (over five-fold) than ciliated cells (p<0.001), but this effect was only moderate on 5 kPa and non-significant for 50 kPa gels (p>0.05). In ciliated cells only, we further examined the effect of stiffness on cell height (figure 2H), cell circularity (figure 2I) and nucleus circularity (figure 2J). Cells on 5 kPa gels had a significantly greater cell height compared to the two other stiffnesses (p<0.001) and cells on all stiffnesses showed transient height changes during serum starvation in all stiffnesses (Figure 2H). All cells showed a non-circular morphology with a mean circularity of less than 0.5. Interestingly, cell circularity but not nucleus circularity was influenced by stiffness. Cells on 5 kPa became significantly more rounded with time (p<0.001), whereas circularity on 50 kPa and coverslips showed only moderate changes. For nuclei circularity, no significant difference was observed across stiffness at all time points (p>0.05).

### Traction force inhibition dysregulated ciliogenesis and cilia characteristics

Cells were next treated with blebbistatin to determine the effect of actomyosin traction forces on ciliogenesis, cilia length and centriole position. Following 24 h of serum starvation, 50 μM blebbistatin was introduced for a further 24 h prior to fixation. Chondrocytes cultured on all substrates appeared more dendritic compared to untreated cells (figure 3A). Blebbistatin treatment significantly reduced mean cilia frequency from 62% to 16% and 59% to 17% in 5 kPa and coverslips, respectively (p<0.001 for both substrates; figure 3B), whereas cilia frequency remained unchanged for cells on 50 kPa gels (p=0.100). In contrast, blebbistatin significantly reduced cilia length in cells cultured on all three substrates (p<0.05; figure 3C). Centriole position (% height) was only significantly affected by blebbistatin in cells on 50 kPa gels, where blebbistatin resulted in a wider distribution of centrioles from the central plane (48% of cell population) compared to untreated controls (p=0.016; figure 3D).

**Figure 3.**
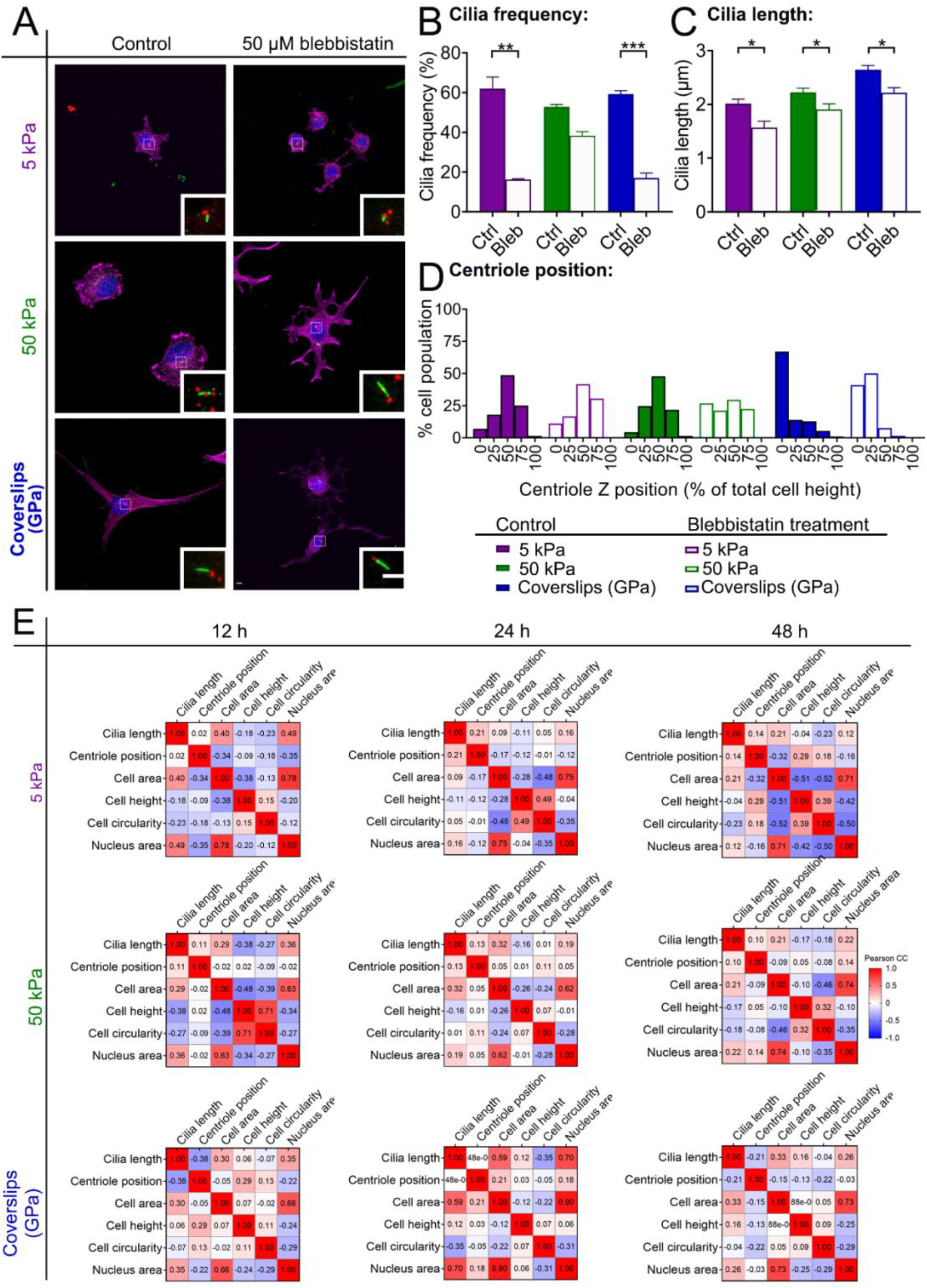
(A) Representative confocal images of control and blebbistatin-treated chondrocytes on different stiffness, stained for primary cilia (ARL13B: green), centriole (γ-tubulin: red), actin (phalloidin: magenta) and nucleus (DAPI: blue). White dashed boxes show the presence of primary cilia. Insets show a magnified view of primary cilia and centriole. Scale bar = 5 μm. Transient changes in (B) cilia frequency (C) cilia length and (D) centriole position after blebbistatin treatment. Ctrl = control, Bleb = blebbistatin treatment. N=3; n ≥66 per condition from a total number of 408 cells. *p<0.05, **p<0.01, ***p<0.001. (E) Correlations between parameters at different stiffness and time points. The correlation was quantified by a Pearson correlation coefficient (r), which is shown in each cell of the correlation matrix. The colour-coded cells provide a direction—red is positive, blue is negative. The parameters were measured from 690 ciliated cells.

### Ciliogenesis, centriole position and cell morphology are dynamically regulated by stiffness

Since we observed relationships between cilia length, cell morphology, and substrate stiffness, we computed pairwise correlations across these parameters. Two representations of correlation are depicted: a heatmap in which colour-coded cells show a Pearson correlation coefficient (r) between two parameters (figure 3E) and a scatter plot with linear regression (figure S2-S4). A time-dependent correlation was observed between cilia length and cell area. A significant positive correlation was observed only at 12 h when cells were cultured on 5 kPa gels but were observed at both 12 h and 24 h when cells were cultured on 50 kPa gels. Remarkably, a significant correlation was observed at all time points of cells cultured on coverslips. The strongest positive correlation was observed between cell and nuclei area, which was consistent at all time points for all stiffnesses (p<0.001). Another interesting finding is how the negative correlation among cell morphology parameters (cell area, cell height and cell circularity) also changed dynamically. At 12 h, the morphological parameters on 5 kPa gels were not correlated with were highly significant by 48 h (a list of all r and p-value is available in supplementary data). Conversely, the opposite was observed for cells on 50 kPa gels. The correlation at 24 h and 48 h was weaker compared to 12 h. Cells on coverslips showed no correlation across cell morphometric parameters over time.

## Discussion

At the tissue level, a key characteristic of osteoarthritis is a softening and eventual destruction of the articular cartilage extracellular matrix. Here, we show that the chondrocyte primary cilium is acutely sensitive to extracellular stiffness, which may have a significant impact on cellular mechanosensation, signal integration and osteoarthritis pathogenesis. In this study, we used substrate stiffness, a known regulator of actin network formation and organisation, to examine the role of substrate stiffness on ciliogenesis in articular chondrocytes. This study has highlighted that the presentation of specific stiffness, with no perturbation of chemical or other physical cell culture cues, can affect primary ciliogenesis and key cell morphology parameters in a stiffness and time-specific manner. In contrast to the current paradigm, we have demonstrated for chondrocytes that stiffer substrates are associated with longer cilia positioned near the cell’s basolateral surface, and that cilia length positively correlates with larger cell areas and increased F-actin distribution.

Substrate stiffness modulation of cilia length is a likely mechanism that enhances chondrocyte mechanosensitivity [38] and it is postulated that an expanded cilia surface area allows for volume signal exposure and signal amplification [39]. The typical length of primary cilium varies among various cell types ranging between 1-10 μm in healthy tissues, and up to 3 μm in chondrocytes [20], [40], [41]. Zhong [42] demonstrated an increase of YAP (Yes-Associated Protein) expression, a key mediator in mechanosignalling, to be significantly elevated when chondrocytes were cultured on a stiffer substrate which supports the idea of enhanced mechanosignalling in longer chondrocyte cilia. The fact that cilia length regulates cell responses to other mechanical stimuli such as compression [12], [13] supports the concept that cilia length is a key effector on the changing substrate stiffness.

Signal transduction in primary cilia is facilitated by the temporal localisation of the associated receptors and protein trafficking [43], [44]. This allows cells to adapt to specific homeostatic cues. In kidney epithelial cells [45], 4 h of fluid flow increase in cilia length, but prolonged exposure (four days) reduced cilia length. Chondrocyte primary cilia are also sensitive to the magnitude and duration of applied load, where prolonged continuous cyclic compression significantly reduced cilia frequency and length, a process that was reversible with stimulus removal. [14]. Our study revealed the importance of timing. Our data demonstrate a highly dynamic process of ciliogenesis and a positive correlation between cilia length and cell area was sustained for longer when cells were cultured on a stiffer substrate. This provides an opportunity to further manipulate cell and cilia biology in a mechanical manner.

We observed no differences in cilia frequency of chondrocytes cultured on different stiffnesses, despite significant changes in the cell area, shape and nuclear morphology. These findings were contradictory to studies in different cell types [46], [47] and could be due to cell lineage and/or cell density. Ciliogenesis is strongly influenced by cell-cell contact, where a higher frequency of ciliogenesis is associated with confluence and reduced mitosis [46], [47]. Further investigations using adhesive micropatterns of increasing size to control individual cell spreading have revealed that instead of cell-cell contact, cell spreading is the major negative influence on ciliogenesis [46], [48]. In keeping with the cartilage phenotype where chondrocytes are separated by several microns of ECM and not to confound the effects on cell crowding on ciliogenesis and actin distribution, we seeded cells at a low density [49]. Temporally, from cell seeding through to 48 h of serum starvation, the number of ciliated cells varied with time and stiffness but by 48 h of serum starvation, cells on all stiffnesses reached very similar ciliation frequency suggesting that cilia frequency is a stiffness-independent process, most likely overridden by the serum concentration. A non-significant trend indicated that cells on softer substrates were slower to ciliate, a process we show is closely regulated to a ‘threshold’ point of cell attachment, spreading and force generation.

A major finding of this study was the effect of substrate stiffness on centriole (and hence primary cilia) positioning. Unlike in non-adherent cells whose centriole sits at the centre of the cell, centriole positioning in adherent cells is tightly regulated [50]. Centriole position supports cells to sense the shape and size by regulating nuclear positioning, which subsequently determines the distribution of organelles [51]. Our findings support that substrate stiffness in chondrocytes impact the centriole position where centrioles locate towards the basolateral surface on stiffer substrates and correlate with increased cell area, actin stress fibres and greater traction forces [46]. Although we acknowledge that in this study, we use a simple model system, which does not replicate the complex heterogenic 3D ECM architecture of cartilage, our data still suggests that even subtle changes of stiffness, e.g. comparing soft (5 kPa) and stiff substrates (50 kPa), can dramatically influence where a centriole and its accompanying cilium is placed. Centriole and primary cilia positioning is critical in development and disease, with well-established roles in roles in tissue morphogenesis, planar cell polarity, and determination of left-right asymmetry. In cartilage, it is important for the detection of morphogen gradients in limb patterning and bone development [9], [21], [29], [46] [53]. Although chondrocytes have no ‘conventional’ apical-basolateral polarity, studies have demonstrated that chondrocytes display distinct positioning within the adult cartilage, the epiphyseal growth plate and osteoarthritic cell clusters [29], [53]–[55]. We believe that the primary ciliium position plays a key role in directing the secretion of ECM for tissue homeostasis; a process which may be defective during OA onset and progression.

Substrate stiffness is key regulator of cell area and spreading, with reduced cellular traction force generation, smaller cell area and more circular cells on softer substrates vice versa on stiffer surfaces [56]. We also observed similar responses with a concomitant change in actin organisation with prominent stress fibres arrangement. However, the few studies that have examined the effect of stiffness and cilia are in epithelial cells, where large actin contractile bundles in highly extended cells limit primary cilia extension from the centriole [46]. In contrast, we observed longer cilia were associated with more prominent actin. Although cell density can contribute to changes in actin distribution, traction force generation and cilia dynamics, comparable studies in low density cultures show opposing data to what is presented in this study. We believe that this supports the notion that cell-type specific properties and lineage (e.g. epithelial versus mesenchymal lineages) drive differential responses to similar external stimuli.

An unexpected stiffness-specific response was the differences in cell area in cells with detectable cilia and those without. We show that on softer substrates, the presence of a cilium did not influence the degree of cell spreading over time compared to non-ciliated cells. However, on the stiffer substrate (coverslip), the presence of a cilium resulted in a large increase in cell area compared to cells without a cilium. This suggests that above a certain stiffness, the absence of a cilium on any substrate maintains or restricts cell area, and the emergence of a cilium is required to drive actin organisation, sufficient traction force generation and to break cellular symmetry for strong cell attachment on the basolateral surface [57]. These findings are also important in developing an understanding of cellular heterogeneity in cell culture models examining substrate rigidity.

In the literature to date, chemical perturbation of actin-myosin (e.g. with cytochalasin or blebbistatin) or RNAi-mediated knockdown promote cilia elongation [47], [58]–[61]. When we examined chondrocytes following blebbistatin exposure, we found cytoskeletal disruption significantly reduced the proportion of ciliated cells, and cilia that remained were shorter, independent of stiffness. These was not due to ineffective disruption of actin as we observed reduced cell shape and a disorganised actin architecture compared to untreated cells on all stiffnesses. So, although these data implicate the involvement of actin contractility in the fine-tuning of cilia length, our data contradicts data in other cell types suggesting a novel mechanism by which articular chondrocytes overcome the contractility barrier conferred by actin on primary cilia elongation.

Our final key finding is in relation to tissue and context-specific stiffness. Stem cells studies have shown non-monotonic responses to substrate stiffness [62], [63], and our further data support this phenomenon. We found that cells on 5 kPa gels and coverslips often shared similar responses despite being cultured on stiffness spanning an order of magnitude. Cilia length on 5 kPa gels and coverslips did not fluctuate significantly over time but did in cells on 50 kPa gels. Our data indicate that the cells confer sensitivity to a specific range of stiffness in a parabolic manner, and this was maintained across various morphological parameters in both ciliated and non-ciliated cells. These findings highlight the importance of culturing cells on or within substrates/ matrices that are relevant to the tissue of origin and disease context. The 5 kPa and 50 kPa stiffness in this study represented osteoarthritic and healthy tissue stiffness, respectively, and our data showed that cilia are shorter on 5 kPa. This mirrors observations in OA chondrocytes [29]. We predict that primary cilia activate different signalling pathways in different stiffness ranges, suggesting that signalling via cilia is sequential rather than constitutive [64]. Our findings show that cells presented with a disease-type stiffness show distinct physiological responses compared to healthy cells, which possibly serve as the origin of disease progression.

Taken together, our data has identified the dynamic interplay between stiffness mechanosignalling and primary cilia length and positioning, and highlights that despite our in-depth appreciation of how primary cilia are such evolutionary conserved organelles, it is critical to understand environmental cues in a mechanically tissue-specific context. This will provide a greater understanding of cilia mechanobiology and how primary cilia are novel targets for mechanotherapy.

## Materials and Methods

PA hydrogels were fabricated as detailed by Tse and Engler [65]. Following hydrogel characterised, chondrocytes were cultured on collagen type II coated hydrogels. After 24 h, chondrocytes were starved in media containing 0.25% FBS for 48 h. Cells were fixed every 12 h. Immunostaining and confocal microscopy were used to characterise primary cilia, cell morphology and nuclei morphology. Correlations between parameters were then calculated for every pair of datasets. To investigate the role of the actomyosin traction force, cells were treated with 50 μM blebbistatin after 24 h in low-serum conditions. Full methods on PA gel fabrication, cell culture, immunostaining, imaging and statistical analysis can be found in SI Materials and Methods.

## Author Contributions

IW, SL, YSC and SRM designed research; IW and SL performed research; AC contributed new reagents/analytic tools; IW, SL and YSC analysed data; IW, SL, YSC, SRM wrote the paper; and YSC and SRM provided financial support. All authors discussed the results and contributed to the final manuscript.

## Acknowledgments

We thank Praju Vikas Anekal and Jacqueline Ross from Biomedical Imaging Research Unit, The University of Auckland for their help with microscopy, Joseph Vella for atomic force microscopy, Jenny Malmstrom and her research group for fruitful discussions on the gel system. This work was supported by Royal Society of New Zealand Marsden Fund (awarded to SRM and YSC). and The University of Auckland Doctoral Funds, and the School of Medical Sciences Graduate Student Fund (to IW).

## References

[1] A. Engler, L. Bacakova, C. Newman, A. Hategan, M. Griffin, and D. Discher, “Substrate compliance versus ligand density in cell on gel responses,” Biophys. J., vol. 86, no. 1, pp. 617–628, 2004.

[2] M. Guo et al., “Cell volume change through water efflux impacts cell stiffness and stem cell fate,” Proc. Natl. Acad. Sci., vol. 114, no. 41, p. E8618 LP–E8627, Oct. 2017.

[3] C.-M. Lo, H.-B. Wang, M. Dembo, and Y. Wang, “Cell Movement Is Guided by the Rigidity of the Substrate,” Biophys. J., vol. 79, no. 1, pp. 144–152, 2000.

[4] A. J. Engler, S. Sen, H. L. Sweeney, and D. E. Discher, “Matrix Elasticity Directs Stem Cell Lineage Specification,” Cell, vol. 126, no. 4, pp. 677–689, 2006.

[5] S. Gobaa, S. Hoehnel, and M. P. Lutolf, “Substrate elasticity modulates the responsiveness of mesenchymal stem cells to commitment cues,” Integr. Biol., vol. 7, no. 10, pp. 1135–1142, 2015.

[6] M. Stolz et al., “Early detection of aging cartilage and osteoarthritis in mice and patient samples using atomic force microscopy,” Nat. Nanotechnol., vol. 4, no. 3, pp. 186–192, 2009.

[7] F. Guilak et al., “The pericellular matrix as a transducer of biomechanical and biochemical signals in articular cartilage,” Ann. N. Y. Acad. Sci., vol. 1068, no. 1, pp. 498–512, 2006.

[8] F. Guilak, L. G. Alexopoulos, M. A. Haider, H. P. Ting-Beall, and L. A. Setton, “Zonal uniformity in mechanical properties of the chondrocyte pericellular matrix: Micropipette aspiration of canine chondrons isolated by cartilage homogenization,” Ann. Biomed. Eng., vol. 33, no. 10, pp. 1312–1318, 2005.

[9] J. J. Malicki and C. A. Johnson, “The Cilium: Cellular Antenna and Central Processing Unit,” Trends Cell Biol., vol. 27, no. 2, pp. 126–140, 2017.

[10] C. A. Poole, M. H. Flint, and B. W. Beaumont, “Analysis of the morphology and function of primary cilia in connective tissues: a cellular cybernetic probe?,” Cell Motil., vol. 5, no. 3, pp. 175–193, 1985.

[11] C. G. Jensen et al., “Ultrastructural, tomographic and confocal imaging of the chondrocyte primary cilium in situ,” Cell Biol. Int., vol. 28, no. 2, pp. 101–110, 2004.

[12] S. R. McGlashan, C. G. Jensen, and C. A. Poole, “Localization of extracellular matrix receptors on the chondrocyte primary cilium,” J. Histochem. Cytochem., vol. 54, no. 9, pp. 1005–1014, 2006.

[13] M. M. Knight, S. R. McGlashan, M. Garcia, C. G. Jensen, and C. A. Poole, “Articular chondrocytes express connexin 43 hemichannels and P2 receptors--a putative mechanoreceptor complex involving the primary cilium?,” J. Anat., vol. 214, no. 2, pp. 275–283, 2009.

[14] S. R. McGlashan et al., “Mechanical loading modulates chondrocyte primary cilia incidence and length.,” Cell Biol. Int., vol. 34, no. 5, pp. 441–446, Mar. 2010.

[15] A. K. T. Wann et al., “Primary cilia mediate mechanotransduction through control of ATP-induced Ca2+ signaling in compressed chondrocytes,” FASEB J., vol. 26, no. 4, pp. 1663–1671, 2012.

[16] J. Irianto, G. Ramaswamy, R. Serra, and M. M. Knight, “Depletion of chondrocyte primary cilia reduces the compressive modulus of articular cartilage,” J. Biomech., vol. 47, no. 2, pp. 579–582, Jan. 2014.

[17] C. L. Thompson, J. P. Chapple, and M. M. Knight, “Primary cilia disassembly down-regulates mechanosensitive hedgehog signalling: a feedback mechanism controlling ADAMTS-5 expression in chondrocytes,” Osteoarthr. Cartil., vol. 22, no. 3, pp. 490–498, 2014.

[18] D. Rux et al., “Primary cilia drive postnatal tidemark patterning in articular cartilage by coordinating responses to Indian Hedgehog and mechanical load,” bioRxiv, 2021.

[19] C. G. Jensen, L. C. W. Jensen, and C. L. Rieder, “The occurrence and structure of primary cilia in a subline of Potorous tridactylus,” Exp. Cell Res., vol. 123, no. 2, pp. 444–449, 1979.

[20] T. Y. Besschetnova, E. Kolpakova-Hart, Y. Guan, J. Zhou, B. R. Olsen, and J. V. Shah, “Identification of Signaling Pathways Regulating Primary Cilium Length and Flow-Mediated Adaptation,” Curr. Biol., vol. 20, no. 2, pp. 182–187, 2010.

[21] R. T. Sherpa, K. F. Atkinson, V. P. Ferreira, and S. M. Nauli, “Rapamycin increases length and mechanosensory function of primary cilia in renal epithelial and vascular endothelial cells,” Int. Educ. Res. J., vol. 2, no. 12, p. 91, 2016.

[22] J. Keeling, L. Tsiokas, and D. Maskey, “Cellular Mechanisms of Ciliary Length Control,” Cells, vol. 5, no. 1, p. 6, 2016.

[23] M.-G. Ascenzi, M. Lenox, and C. Farnum, “Analysis of the orientation of primary cilia in growth plate cartilage: A mathematical method based on multiphoton microscopical images,” J. Struct. Biol., vol. 158, no. 3, pp. 293–306, 2007.

[24] C. E. Farnum and N. J. Wilsman, “Axonemal positioning and orientation in three-dimensional space for primary cilia: What is known, what is assumed, and what needs clarification,” Dev. Dyn., vol. 240, no. 11, pp. 2405–2431, 2011.

[25] H. A. Praetorius and K. R. Spring, “Removal of the MDCK cell primary cilium abolishes flow sensing,” J. Membr. Biol., vol. 191, no. 1, pp. 69–76, 2003.

[26] C. J. Haycraft et al., “Intraflagellar transport is essential for endochondral bone formation,” Development, vol. 134, no. 2, pp. 307–316, Jan. 2007.

[27] S. R. McGlashan, C. J. Haycraft, C. G. Jensen, B. K. Yoder, and C. A. Poole, “Articular cartilage and growth plate defects are associated with chondrocyte cytoskeletal abnormalities in Tg737orpk mice lacking the primary cilia protein polaris,” Matrix Biol., vol. 26, no. 4, pp. 234–246, 2007.

[28] B. Song, C. J. Haycraft, H. Seo, B. K. Yoder, and R. Serra, “Development of the post-natal growth plate requires intraflagellar transport proteins,” Dev. Biol., vol. 305, no. 1, pp. 202–216, 2007.

[29] S. R. McGlashan, E. C. Cluett, C. G. Jensen, and C. A. Poole, “Primary Cilia in osteoarthritic chondrocytes: From chondrons to clusters,” Dev. Dyn., vol. 237, no. 8, pp. 2013–2020, 2008.

[30] Y. S. Choi et al., “The alignment and fusion assembly of adipose-derived stem cells on mechanically patterned matrices,” Biomaterials, vol. 33, no. 29, pp. 6943–6951, 2012.

[31] S. Lelièvre, V. M. Weaver, and M. J. Bissell, “Extracellular matrix signaling from the cellular membrane skeleton to the nuclear skeleton: a model of gene regulation,” Recent Prog. Horm. Res., vol. 51, p. 417, 1996.

[32] R. McBeath, D. M. Pirone, C. M. Nelson, K. Bhadriraju, and C. S. Chen, “Cell shape, cytoskeletal tension, and RhoA regulate stem cell lineage commitment,” Dev. Cell, vol. 6, no. 4, pp. 483–495, 2004.

[33] C. C. DuFort, M. J. Paszek, and V. M. Weaver, “Balancing forces: architectural control of mechanotransduction.,” Nat. Rev. Mol. Cell Biol., vol. 12, no. 5, pp. 308–319, 2011.

[34] J. K. Mouw, G. Ou, and V. M. Weaver, “Extracellular matrix assembly: a multiscale deconstruction,” Nat. Rev. Mol. cell Biol., vol. 15, no. 12, pp. 771–785, 2014.

[35] J. Kim et al., “Functional genomic screen for modulators of ciliogenesis and cilium length,” Nature, vol. 464, no. 7291, pp. 1048–1051, 2010.

[36] A. Karim and A. C. Hall, “Chondrocyte Morphology in Stiff and Soft Agarose Gels and the Influence of Fetal Calf Serum,” J. Cell. Physiol., vol. 232, no. 5, pp. 1041–1052, May 2017.

[37] E. Schuh et al., “Effect of Matrix Elasticity on the Maintenance of the Chondrogenic Phenotype,” Tissue Eng. Part A, vol. 16, no. 4, pp. 1281–1290, Nov. 2009.

[38] M. Spasic and C. R. Jacobs, “Lengthening primary cilia enhances cellular mechanosensitivity,” Eur. Cell. Mater., vol. 33, pp. 158–168, Feb. 2017.

[39] D. K. Breslow, E. F. Koslover, F. Seydel, A. J. Spakowitz, and M. V Nachury, “An in vitro assay for entry into cilia reveals unique properties of the soluble diffusion barrier,” J. Cell Biol., vol. 203, no. 1, pp. 129–147, Oct. 2013.

[40] J. R. Broekhuis, S. Rademakers, J. Burghoorn, and G. Jansen, “SQL-1, homologue of the Golgi protein GMAP210, modulates intraflagellar transport in *gt;C. elegans*,” J. Cell Sci., vol. 126, no. 8, p. 1785 LP–1795, Apr. 2013.

[41] R. M. Delaine-Smith, A. Sittichokechaiwut, and G. C. Reilly, “Primary cilia 518 respond to fluid shear stress and mediate flow-induced calcium deposition in 519 osteoblasts,” FASEB J. Off. Publ. Fed. Am. 520 Soc. Exp. Biol., vol. 28, no. 430–439, p. 521, 2014.

[42] W. Zhong, Y. Li, L. Li, W. Zhang, S. Wang, and X. Zheng, “YAP-mediated regulation of the chondrogenic phenotype in response to matrix elasticity,” J. Mol. Histol., vol. 44, no. 5, pp. 587–595, Oct. 2013.

[43] Z. Anvarian, K. Mykytyn, S. Mukhopadhyay, L. B. Pedersen, and S. T. Christensen, “Cellular signalling by primary cilia in development, organ function and disease,” Nat. Rev. Nephrol., vol. 15, no. 4, pp. 199–219, Apr. 2019.

[44] M. V Nachury, “The molecular machines that traffic signaling receptors into and out of cilia.,” Curr. Opin. Cell Biol., vol. 51, pp. 124–131, Apr. 2018.

[45] I. Orhon et al., “Primary-cilium-dependent autophagy controls epithelial cell volume in response to fluid flow,” Nat. Cell Biol., vol. 18, no. 6, pp. 657–667, 2016.

[46] A. Pitaval, Q. Tseng, M. Bornens, and M. Théry, “Cell shape and contractility regulate ciliogenesis in cell cycle-arrested cells,” J. Cell Biol., vol. 191, no. 2, pp. 303–312, 2010.

[47] J. Kim et al., “Actin remodelling factors control ciliogenesis by regulating YAP/TAZ activity and vesicle trafficking,” Nat. Commun., vol. 6, pp. 1–13, 2015.

[48] T. Yeung et al., “Effects of substrate stiffness on cell morphology, cytoskeletal structure, and adhesion,” Cell Motil. Cytoskeleton, vol. 60, no. 1, pp. 24–34, 2005.

[49] A. C. Hall, “The Role of Chondrocyte Morphology and Volume in Controlling Phenotype—Implications for Osteoarthritis, Cartilage Repair, and Cartilage Engineering,” Curr. Rheumatol. Rep., vol. 21, no. 8, p. 38, 2019.

[50] M. Théry et al., “Anisotropy of cell adhesive microenvironment governs cell internal organization and orientation of polarity,” Proc. Natl. Acad. Sci., vol. 103, no. 52, p. 19771 LP–19776, Dec. 2006.

[51] J. Zhu, A. Burakov, V. Rodionov, and A. Mogilner, “Finding the Cell Center by a Balance of Dynein and Myosin Pulling and Microtubule Pushing: A Computational Study,” Mol. Biol. Cell, vol. 21, no. 24, pp. 4418–4427, Oct. 2010.

[52] T. J. Park, S. L. Haigo, and J. B. Wallingford, “Ciliogenesis defects in embryos lacking inturned or fuzzy function are associated with failure of planar cell polarity and Hedgehog signaling,” Nat. Genet., vol. 38, no. 3, pp. 303–311, 2006.

[53] C. E. de Andrea, M. Wiweger, F. Prins, J. V. M. G. Bovée, S. Romeo, and P. C. W. Hogendoorn, “Primary cilia organization reflects polarity in the growth plate and implies loss of polarity and mosaicism in osteochondroma,” Lab. Investig., vol. 90, no. 7, pp. 1091–1101, 2010.

[54] C. T. G. Appleton, V. Pitelka, J. Henry, and F. Beier, “Global analyses of gene expression in early experimental osteoarthritis,” Arthritis Rheum., vol. 56, no. 6, pp. 1854–1868, Jun. 2007.

[55] E. V Tchetina, G. Squires, and A. R. Poole, “Increased type II collagen degradation and very early focal cartilage degeneration is associated with upregulation of chondrocyte differentiation related genes in early human articular cartilage lesions.,” J. Rheumatol., vol. 32, no. 5, p. 876 LP–886, May 2005.

[56] J. Escribano, M. T. Sánchez, and J. M. García-Aznar, “A discrete approach for modeling cell–matrix adhesions,” Comput. Part. Mech., vol. 1, no. 2, pp. 117–130, 2014.

[57] B. L. Doss et al., “Cell response to substrate rigidity is regulated by active and passive cytoskeletal stress,” Proc. Natl. Acad. Sci., vol. 117, no. 23, p. 12817 LP–12825, Jun. 2020.

[58] J. Cao et al., “MiR-129-3p controls cilia assembly by regulating CP110 and actin dynamics,” Nat. Cell Biol., vol. 14, no. 7, pp. 697–706, 2012.

[59] A. Li et al., “Ciliary transition zone activation of phosphorylated Tctex-1 controls ciliary resorption, S-phase entry and fate of neural progenitors,” Nat. Cell Biol., vol. 13, no. 4, pp. 402–411, 2011.

[60] S. Tomoshige, Y. Kobayashi, K. Hosoba, A. Hamamoto, T. Miyamoto, and Y. Saito, “Cytoskeleton-related regulation of primary cilia shortening mediated by melanin-concentrating hormone receptor 1,” Gen. Comp. Endocrinol., vol. 253, pp. 44–52, 2017.

[61] P. Kohli et al., “The ciliary membrane-associated proteome reveals actin-binding proteins as key components of cilia,” EMBO Rep., vol. 18, no. 9, p. e201643846, 2017.

[62] J. Swift et al., “Nuclear Lamin-A Scales with Tissue Stiffness and Enhances Matrix-Directed Differentiation,” Science (80-.)., vol. 341, no. 6149, p. 1240104, Aug. 2013.

[63] C. Kim et al., “Stem Cell Mechanosensation on Gelatin Methacryloyl (GelMA) Stiffness Gradient Hydrogels,” Ann. Biomed. Eng., vol. 48, no. 2, pp. 893–902, 2020.

[64] R. Pala, N. Alomari, and S. M. Nauli, “Primary Cilium-Dependent Signaling Mechanisms,” Int. J. Mol. Sci., vol. 18, no. 11, p. 2272, Oct. 2017.

[65] J. R. Tse and A. J. Engler, “Preparation of Hydrogel Substrates with Tunable Mechanical Properties,” Curr. Protoc. Cell Biol., vol. 47, no. 1, p. 10.16.1–10.16.16, Jun. 2010.

