## Supplementary information for "Matrix stiffness controls ciliogenesis and centriole position"

### Supplementary Information for
Matrix stiffness controls ciliogenesis and centriole position

Ivanna Williantarra[1], Sophia Leung[1], Yu Suk Choi[2], Ashika Channa[1], Sue R McGlashan[1]*
1 Department of Anatomy and Medical Imaging, Faculty of Medical and Health Sciences, University of Auckland, New Zealand
2 School of Human Sciences, University of Western Australia, Australia

*Sue R McGlashan


**This PDF file includes:**

Supplementary materials and methods
Figures S1 to S4
Tables S1 to S9

#### Supplementary Materials and Methods

##### PA gel fabrication

Glass coverslips (#1,ProSciTech,Australia) were first sterilized by UV radiation (UV/Ozone ProCleanerTM, BioForce Nanosciences, United States of America) and then functionalized using 3% acetic acid (Scharlau, Spain) and 0.5% 3-(trimethoxysilyl)propylmethacrylate (Sigma Aldrich, New Zealand) in 100% ethanol (Merck, New Zealand) to facilitate hydrogel covalent attachment. 5 kPa and 50 kPa hydrogels were produced by mixing 40% w/v acrylamide monomer and 2% w/v N,N methylene-bis-acrylamide crosslinker (Bio-Rad, New Zealand) in a ratio of 10:1 and 30:1, respectively. The polymerization was induced by addition of 0.1% w/v degassed ammonium persulfate (APS; Bio-Rad New Zealand) and 0.1% N,N,N',N'-tetramethylethylenediamine (Bio-Rad, New Zealand). The polymerizing solution was sandwiched between a functionalized coverslip and a dichlorodimethylsilane-treated slide (Sigma Aldrich, New Zealand) to ensure easy detachment, and let to polymerize for at least 15 minutes. After polymerization, hydrogels were then transferred to a 6-well plate, hydrated with dPBS and stored at 4°C until further use. One day prior to cell seeding, gels were functionalized with 0.1 mg/ml Sulfo-SANPAH (sulfosuccinimidyl 6-(4'-azido-2'-nitrophenylamino)hexanoate (Life Technologies, New Zealand) that was activated with 15 W ultraviolet light (350 nm; DH-WUV00050, Interlab New Zealand) for 10 minutes. Gels were washed with PBS then incubated overnight in 0.01 mg/ml collagen type II (Sigma Aldrich, New Zealand) for cell adhesion. Collagen type II coated glass coverslips were used as controls. Excess collagen solution was removed and gels and coverslips were washed with PBS before seeding with cells.

##### Atomic Force Microscopy

PA gel elasticity was quantified by nano-indentation using MFP-3D Origin Atomic Force Microscope (AFM Asylum Research Oxford Instruments Group, United States of America). A total of 30 gels per stiffness were fabricated for each batch. Two gels per batch were randomly selected to verify batch stiffness consistency. Surface indentations were performed using a 200 μm gold-coated, silicon nitride triangular-shaped cantilever tips with a 0.8 N/m force constants, 17 kHz resonance frequency, and a pyramidal height of 3.5 μm (Nano World, UnitedStates of America). Samples were indented with an approach velocity of 2 μm/s until a trigger point of 2 nN was registered and retracted at a velocity of 10 μm/s. Starting 1 mm away from the gel edge, gels were indented in triplicate every 2 mm along the midline. The generated force curves were then analyzed using custom-written code in Igor Pro to determine Young's modulus as previously described [86].

##### Cell culture

Immortalized mouse articular chondrocytes (H5 clone; kind gift from P.M. van der Kraan and HM van Beuningen, Radboud University Medical Center, Nijmegen, The Netherlands) were seeded at a density of 40 cells/mm$^2$ on both PA-gels and glass coverslips. Cells were cultured in Dulbecco's Modified Eagle Medium (DMEM) high glucose, high pyruvate (Life Technologies, New Zealand), supplemented with 10% foetal bovine serum (FBS, In Vitro Technologies, New Zealand) and 1% penicillin/streptomycin (Life Technologies, New Zealand) at 37°C and 5% $CO_2$ for 24 hours. After 24 hours following cell seeding, the culture medium was then removed, and samples were rinsed in PBS and cultured in media supplemented with 0.25% FBS for a further 48 hours. Cells were fixed at 12 hour intervals following the introduction of low serum media conditions using 4% paraformaldehyde for 20 minutes at 37$^o$C (PFA, Sigma Aldrich New Zealand).

For blebbistatin treatment, (-)-blebbistatin (Sigma-Aldrich, New Zealand) was dissolved in DMSO to make a 100-mM stock solution. After 24 hours in low-serum condition, cells were treated with 50 μM blebbistatin for another 24 hours, followed by PFA fixation.

##### Immunostaining

Fixed samples were washed three times with PBS, permeabilized with 0.5% (v/v, in PBS) Triton X-100 (Sigma Aldrich, New Zealand) for 5 minutes, then washed in 0.1% w/v bovine serum albumin in PBS (PBS-BSA, Applichem, Spain) and then blocked with 5% normal goat serum (Sigma Aldrich, New Zealand) in PBS-BSA solution for 30 minutes at room temperature. Samples were incubated simultaneously with rabbit polyclonal ARL13B (1:500, Proteintech, United States of America) and mouse monoclonal γ-tubulin (1:500, SigmaAldrich, New Zealand) overnight at 4$^o$C. Samples were then washed 3 times in PBS-BSA solution before incubation with goat anti-rabbit Alexa 594 and goat anti-mouse Alexa 647 (1:1000, Life Technologies, New Zealand) for 1 hour at room temperature, and then washed three times in PBS-BSA solution. To label filamentous actin (F-actin), samples were incubated with Alexa 488 Phalloidin (1:40, Life Technologies, New Zealand) for 20 minutes at room temperature. Samples were washed three times in PBS-BSA solution before mounting onto slides with ProLong Gold antifading reagent containing DAPI (Life Technologies, New Zealand).

**Image Acquisition and Processing**

Acquired images were used to measure cilia characteristics and cell morphometrics. Cilia characteristics included cilia frequency, cilia length, centriole position and cilia orientation. Cell morphology included cell area, cell height, cell circularity, nucleus area and nucleus circularity. Cilia frequency was examined every 12 hours, starting 12 hours before starvation until 48 hours after starvation, while all other parameters were examined at 12, 24 and 48 hours after starvation.

To quantify cilia frequency, samples were imaged using a Leica DMR upright fluorescence microscope with a 100x/1.4 NA (Numerical Aperture) oil objective. Cilia frequency was presented as the percentage of ciliated cells in a cell population, with a minimum number of 100 cells for each sample (n=300 cells per condition from N=3 replicates).

For other parameters, a minimum of 20 cells was sampled for each condition (n>60 cells per condition from N=3 replicates). Images were acquired using Olympus FV-1000 confocal microscope (Biomedical Imaging Research Unit, The University of Auckland) equipped with a UPLSAPO 100x/1.4 NA objective (Olympus Life Science) and a fully motorized BX61 stage. The confocal aperture was set to 195 μm. Four channels were used to detect DAPI, Alexa 488, Alexa 594 and Alexa 647 fluorophores. Images were acquired using FluoView 4.2 software. Serial optical z-sections were collected with a pixel dwell time of 4 μs/pixel. Up to 30 optical confocal sections spaced 0.5 μm apart encompassing the entire height of cells were acquired. Each image was 1024 x 1024 pixels with a voxel size of 0.215 μm x 0.215 μm x 0.943 μm.

Serial optical z sections were used to create 2D maximum z-projections to measure cilia length, cell area (for both ciliated and non-ciliated cells), cell circularity, nucleus area and nucleus circularity. XZ-projections were used to measure centriole position, cilia orientation and cell height. All parameters were measured using Fiji ImageJ software (https://imagej.nih.gov/ij/).

To determine cilia length, the line tool in Fiji ImageJ was used to measure the distance between the proximal and distal end of the primary cilium from the maximum intensity projection of confocal z-stack images. To measure cell and nucleus area, image contrast was first adjusted and then segmented using the threshold command. Cells or nuclei were selected using the versatile wand tool. Area and circularity were then measured using the area and shape descriptor command. Circularity is presented as a value between 0 to 1. A value approaching 0 indicates an increasingly elongated polygon, while a value of 1 indicates a perfect circle.

XZ projections were generated using XYZ projection tool in Fiji ImageJ. Cell height was measured by drawing a line centred between the highest part of the apical side and basal sides of the cell. Cilia position was expressed as the percentage ratio between centriole z-position and individual cell height (figure 1E). Centrioles were classified to be positioned at the cell's apical domain when positioned between 88%-100% of total cell height, subapical domain when positioned between 63%-87%, central domain between 38%-62% and the rest is the basal domain. For cilia orientation, the elevation angle (Φ) of primary cilia respect to the XY plane was measured by drawing a line from cilia proximal to the distal end of their XZ projection images. The distribution of the elevation angle ranges from -90° to 90° ($-90^o \leq \Phi \leq 90^o$).

**Statistics**

All data are presented as the mean ± standard error of the mean (SEM). Comparisons between means of two groups were performed using an unpaired Student's t-test using Graphpad Prism 8 (https://www.graphpad.com/scientific-software/prism/). When 2 variables were involved, a two-way ANOVA test was used, and a significant interaction was interpreted using a post-hoc Tukey's or Bonferroni's multiple comparisons test. A p-value less than 5% was deemed statistically significant. P values (p) are represented as follows: * = p<0.05, ** = p<0.01, *** = p<0.001.

A heatmap correlation matrix was constructed by calculating the significance of Pearson's correlation coefficient, r, for every pair of datasets using Graphpad Prism 8. Data are assumed to follow Gaussian distribution with 95% confidence interval.

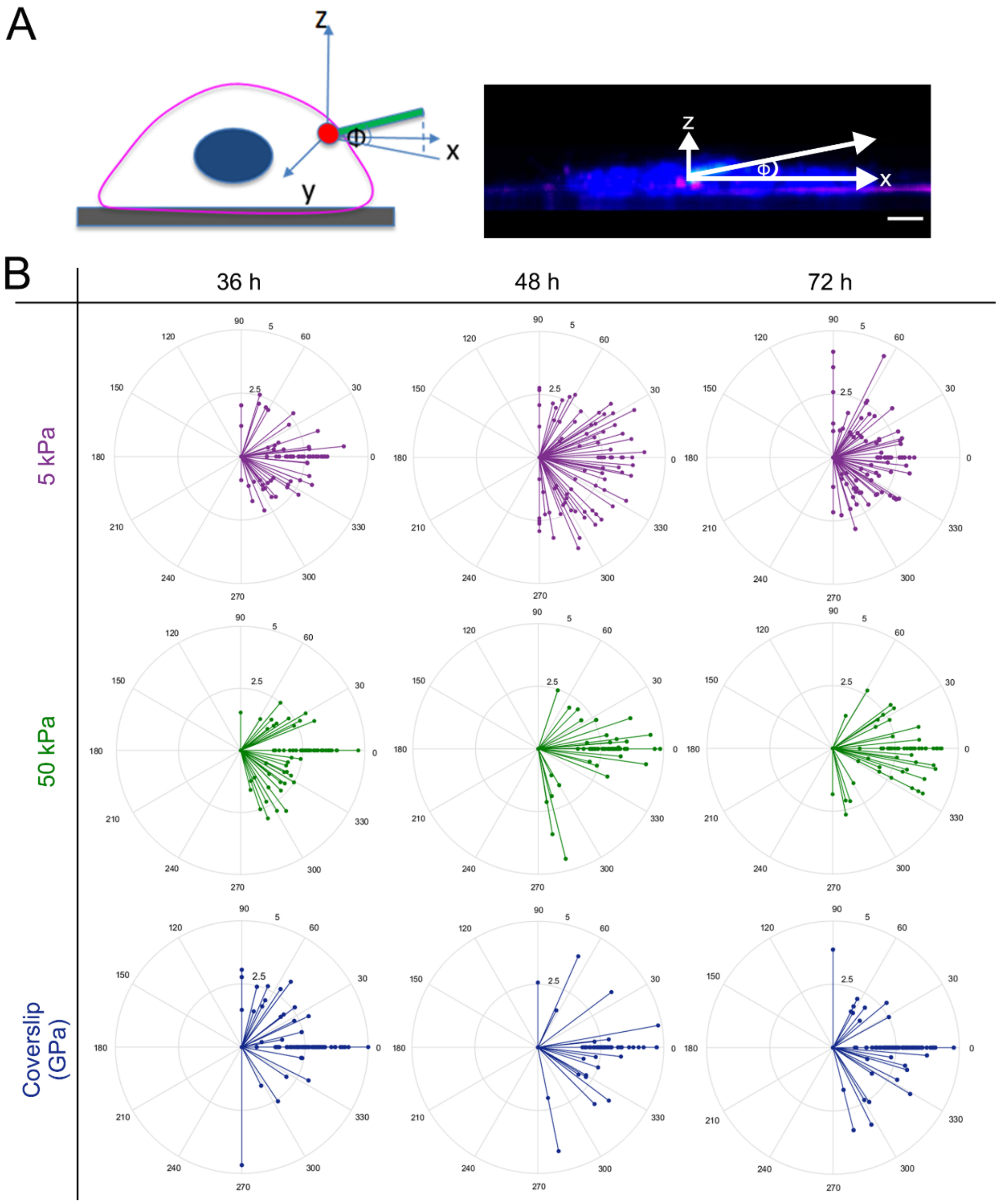


**Fig. S1.** (A) Schematic of a chondrocyte primary cilium showing ciliary elevation angle Φ. Scale bar = 5 μm. (B) Distributions of ciliary elevation angle Φ (-90°≤Φ≤90°) over time. No significant difference in cilia orientation was observed across stiffness. The length of the lines indicates the primary cilium's length. N = 3; n, ≥66 per stiffness. A total number of 690 cells were examined.

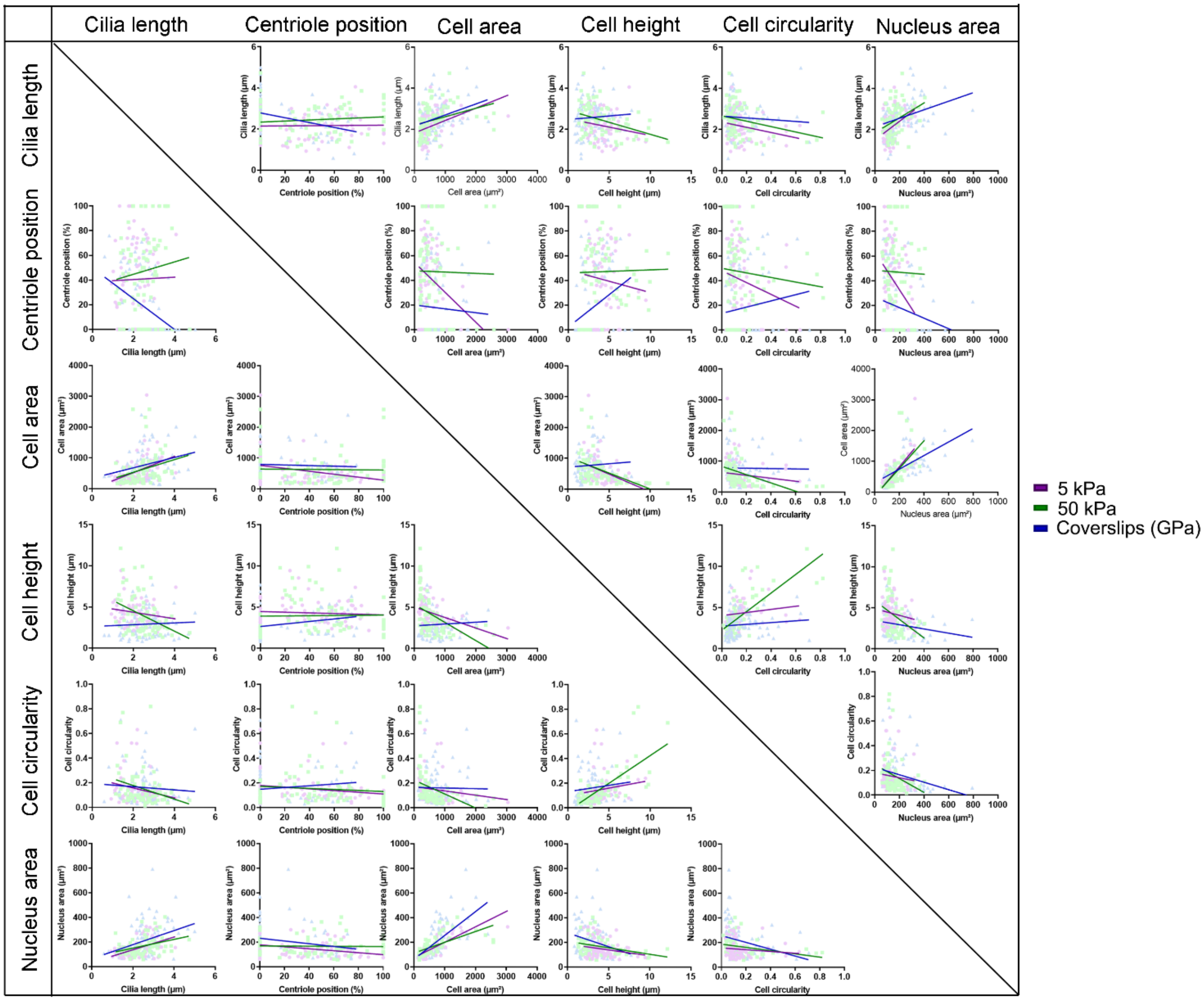


**Fig. S2.** Scatter plot and linear regression between parameters at 12 h

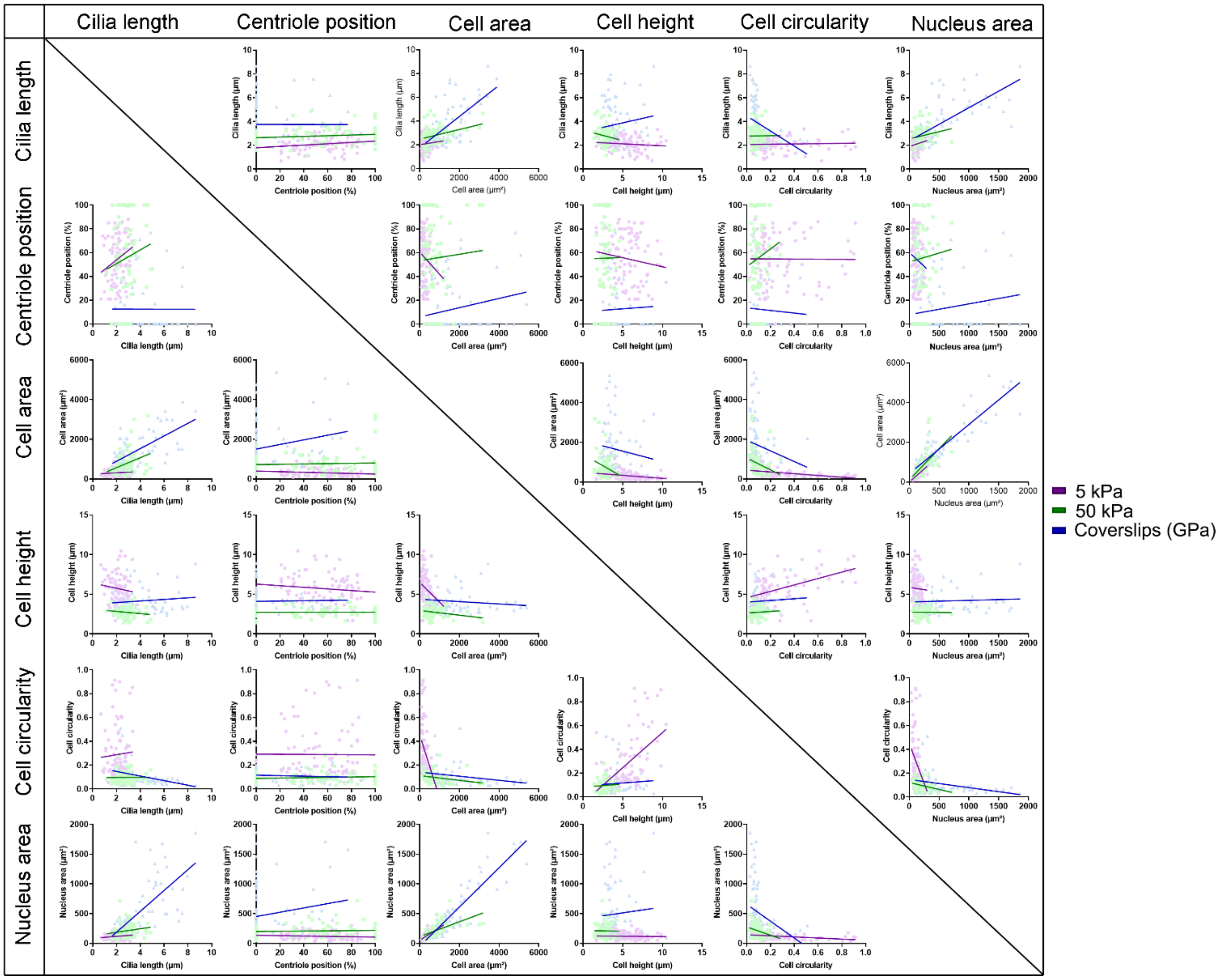


**Fig. S3.** Scatter plot and linear regression between parameters at 24 h

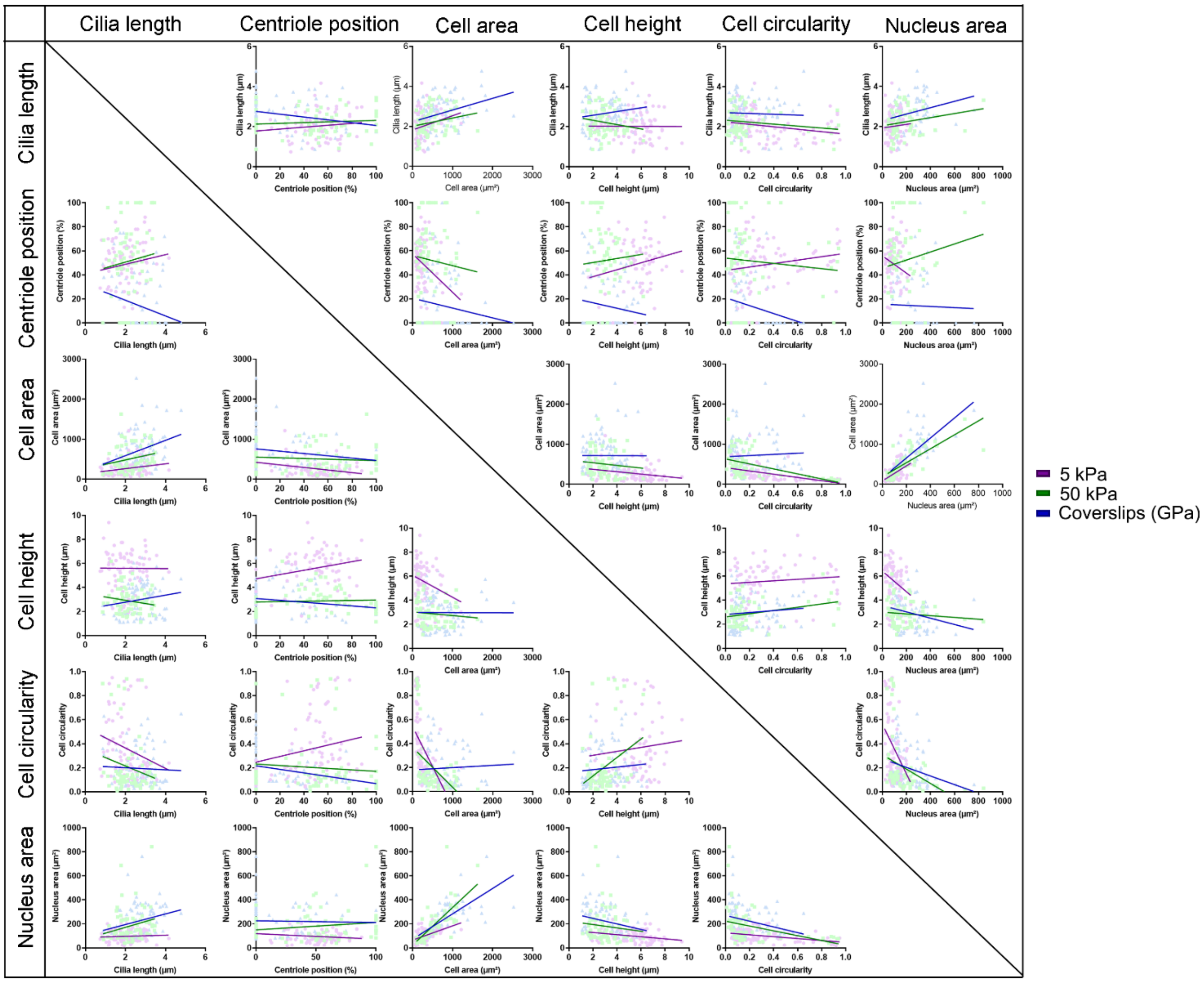


**Fig. S4.** Scatter plot and linear regression between parameters at 24 h

**Table S1.** Correlation test summary of 12 hours, 5 kPa

Summary statistics (Quantitative data):

| Variable | Observations | Minimum | Maximum | Mean | Std. deviation |
|---|---|---|---|---|---|
| Cilia length | 68 | 0.930 | 4.060 | 2.165 | 0.677 |
| Centriole position | 68 | 0.000 | 100.000 | 40.618 | 31.900 |
| Cell area | 68 | 138.050 | 3041.475 | 566.284 | 446.659 |
| Cell height | 68 | 1.938 | 9.440 | 4.299 | 1.492 |
| Cell circularity | 68 | 0.032 | 0.632 | 0.152 | 0.121 |
| Nucleus area | 68 | 61.030 | 325.116 | 145.495 | 48.098 |

Correlation matrix (Pearson):

| Variables | Cilia length | Centriole position | Cell area | Cell height | Cell circularity | Nucleus area |
|---|---|---|---|---|---|---|
| Cilia length | **1** | 0.020 | **0.402** | -0.179 | -0.226 | **0.246** |
| Centriole position | 0.020 | **1** | **-0.340** | -0.086 | -0.181 | **-0.347** |
| Cell area | **0.402** | **-0.340** | **1** | **-0.382** | -0.129 | **0.777** |
| Cell height | -0.179 | -0.086 | **-0.382** | **1** | 0.155 | -0.196 |
| Cell circularity | -0.226 | -0.181 | -0.129 | 0.155 | **1** | -0.119 |
| Nucleus area | **0.246** | **-0.347** | **0.777** | -0.196 | -0.119 | **1** |

*P*-values (Pearson):

| Variables | Cilia length | Centriole position | Cell area | Cell height | Cell circularity | Nucleus area |
|---|---|---|---|---|---|---|
| Cilia length | **0** | 0.869 | **0.001** | 0.145 | 0.064 | **<0.001** |
| Centriole position | 0.869 | **0** | **0.005** | 0.248 | 0.139 | **0.004** |
| Cell area | **0.001** | **0.005** | **0** | **0.001** | 0.294 | **<0.001** |
| Cell height | 0.145 | 0.248 | **0.001** | **0** | 0.208 | 0.110 |
| Cell circularity | 0.064 | 0.139 | 0.294 | 0.208 | **0** | 0.332 |
| Nucleus area | **<0.001** | **0.004** | **<0.001** | 0.110 | 0.332 | **0** |

Coefficients of determination (Pearson):

| Variables | Cilia length | Centriole position | Cell area | Cell height | Cell circularity | Nucleus area |
|---|---|---|---|---|---|---|
| Cilia length | **1** | 0.000 | **0.161** | 0.032 | 0.051 | **0.237** |
| Centriole position | 0.000 | **1** | **0.116** | 0.007 | 0.033 | **0.121** |
| Cell area | **0.161** | **0.116** | **1** | **0.146** | 0.017 | **0.604** |
| Cell height | 0.032 | 0.007 | **0.146** | **1** | 0.024 | 0.038 |
| Cell circularity | 0.051 | 0.033 | 0.017 | 0.024 | **1** | 0.014 |
| Nucleus area | **0.237** | **0.121** | **0.604** | 0.038 | 0.014 | **1** |

Shaded values in bold are different from 0 with a significance level alpha=0.05

**Table S2.** Correlation test summary of 24 hours, 5 kPa

Summary statistics (Quantitative data):

| Variable | Observations | Minimum | Maximum | Mean | Std. deviation |
|---|---|---|---|---|---|
| Cilia length | 82 | 0.690 | 3.128 | 2.101 | 0.601 |
| Centriole position | 82 | 0.000 | 100.000 | 54.695 | 22.696 |
| Cell area | 82 | 85.370 | 1212.111 | 312.165 | 204.708 |
| Cell height | 82 | 1.670 | 10.460 | 5.483 | 1.860 |
| Cell circularity | 82 | 0.029 | 0.913 | 0.289 | 0.225 |
| Nucleus area | 82 | 33.301 | 299.380 | 117.826 | 58.401 |

Correlation matrix (Pearson):

| Variables | Cilia length | Centriole position | Cell area | Cell height | Cell circularity | Nucleus area |
|---|---|---|---|---|---|---|
| Cilia length | **1** | 0.213 | 0.092 | -0.106 | 0.045 | 0.163 |
| Centriole position | 0.213 | **1** | -0.168 | -0.125 | -0.006 | -0.121 |
| Cell area | 0.092 | -0.168 | **1** | **-0.275** | **-0.244** | **0.754** |
| Cell height | -0.106 | -0.125 | **-0.275** | **1** | **0.249** | -0.012 |
| Cell circularity | 0.045 | -0.006 | **-0.244** | **0.249** | **1** | **-0.350** |
| Nucleus area | 0.163 | -0.121 | **0.754** | -0.012 | **-0.350** | **1** |

*P*-values (Pearson):

| Variables | Cilia length | Centriole position | Cell area | Cell height | Cell circularity | Nucleus area |
|---|---|---|---|---|---|---|
| Cilia length | **0** | 0.055 | 0.410 | 0.345 | 0.687 | 0.144 |
| Centriole position | 0.055 | **0** | 0.130 | 0.265 | 0.958 | 0.280 |
| Cell area | 0.410 | 0.130 | **0** | **0.012** | **<0.001** | **<0.001** |
| Cell height | 0.345 | 0.265 | **0.012** | **0** | **<0.001** | 0.751 |
| Cell circularity | 0.687 | 0.958 | **<0.001** | **<0.001** | **0** | **0.001** |
| Nucleus area | 0.144 | 0.280 | **<0.001** | 0.751 | **0.001** | **0** |

Coefficients of determination (Pearson):

| Variables | Cilia length | Centriole position | Cell area | Cell height | Cell circularity | Nucleus area |
|---|---|---|---|---|---|---|
| Cilia length | **1** | 0.045 | 0.009 | 0.011 | 0.002 | 0.027 |
| Centriole position | 0.045 | **1** | 0.028 | 0.016 | 0.000 | 0.015 |
| Cell area | 0.009 | 0.028 | **1** | **0.076** | **0.234** | **0.569** |
| Cell height | 0.011 | 0.016 | **0.076** | **1** | **0.239** | 0.001 |
| Cell circularity | 0.002 | 0.000 | **0.234** | **0.239** | **1** | **0.122** |
| Nucleus area | 0.027 | 0.015 | **0.569** | 0.001 | **0.122** | **1** |

*Shaded values in bold are different from 0 with a significance level alpha=0.05*

**Table S3.** Correlation test summary of 48 hours, 5 kPa

Summary statistics (Quantitative data):

| Variable | Observations | Minimum | Maximum | Mean | Std. deviation |
|---|---|---|---|---|---|
| Cilia length | 48 | 0.740 | 4.170 | 2.015 | 0.703 |
| Centriole position | 48 | 0.000 | 88.000 | 24.889 | 20.376 |
| Cell area | 48 | 61.660 | 1214.780 | 265.322 | 204.592 |
| Cell height | 48 | 1.620 | 9.400 | 5.590 | 1.603 |
| Cell circularity | 48 | 0.040 | 0.950 | 0.123 | 0.262 |
| Nucleus area | 48 | 17.890 | 349.230 | 99.381 | 61.225 |

Correlation matrix (Pearson):

| Variables | Cilia length | Centriole position | Cell area | Cell height | Cell circularity | Nucleus area |
|---|---|---|---|---|---|---|
| Cilia length | **1** | 0.138 | 0.208 | -0.041 | -0.228 | 0.120 |
| Centriole position | 0.138 | **1** | **-0.322** | **0.290** | 0.184 | -0.161 |
| Cell area | 0.208 | **-0.322** | **1** | **-0.508** | **-0.521** | **0.711** |
| Cell height | -0.041 | **0.290** | **-0.508** | **1** | **0.387** | **-0.424** |
| Cell circularity | -0.228 | 0.184 | **-0.521** | **0.387** | **1** | **-0.503** |
| Nucleus area | 0.120 | -0.161 | **0.711** | **-0.424** | **-0.503** | **1** |

*P*-values (Pearson):

| Variables | Cilia length | Centriole position | Cell area | Cell height | Cell circularity | Nucleus area |
|---|---|---|---|---|---|---|
| Cilia length | **0** | 0.247 | 0.080 | 0.730 | 0.054 | 0.315 |
| Centriole position | 0.247 | **0** | **0.006** | **0.013** | 0.121 | 0.177 |
| Cell area | 0.080 | **0.006** | **0** | **<0.001** | **<0.001** | **<0.001** |
| Cell height | 0.730 | **0.013** | **<0.001** | **0** | **0.001** | **<0.001** |
| Cell circularity | 0.054 | 0.121 | **<0.001** | **0.001** | **0** | **<0.001** |
| Nucleus area | 0.315 | 0.177 | **<0.001** | **<0.001** | **<0.001** | **0** |

Coefficients of determination (Pearson):

| Variables | Cilia length | Centriole position | Cell area | Cell height | Cell circularity | Nucleus area |
|---|---|---|---|---|---|---|
| Cilia length | **1** | 0.019 | 0.043 | 0.002 | 0.052 | 0.014 |
| Centriole position | 0.019 | **1** | **0.104** | **0.084** | 0.034 | 0.026 |
| Cell area | 0.043 | **0.104** | **1** | **0.258** | **0.271** | **0.505** |
| Cell height | 0.002 | **0.084** | **0.258** | **1** | **0.150** | **0.179** |
| Cell circularity | 0.052 | 0.034 | **0.271** | **0.150** | **1** | **0.253** |
| Nucleus area | 0.014 | 0.026 | **0.505** | **0.179** | **0.253** | **1** |

*Shaded values in bold are different from 0 with a significance level alpha=0.05*

**Table S4.** Correlation test summary of 12 hours, 50 kPa

Summary statistics (Quantitative data):

| Variable | Observations | Minimum | Maximum | Mean | Std. deviation |
|---|---|---|---|---|---|
| Cilia length | 73 | 1.150 | 4.734 | 2.462 | 0.734 |
| Centriole position | 73 | 0.000 | 100.000 | 47.082 | 32.225 |
| Cell area | 73 | 164.220 | 2577.222 | 627.617 | 518.858 |
| Cell height | 73 | 1.300 | 12.140 | 3.966 | 2.395 |
| Cell circularity | 73 | 0.015 | 0.820 | 0.153 | 0.151 |
| Nucleus area | 73 | 57.410 | 403.780 | 166.737 | 48.623 |

Correlation matrix (Pearson):

| Variables | Cilia length | Centriole position | Cell area | Cell height | Cell circularity | Nucleus area |
|---|---|---|---|---|---|---|
| Cilia length | **1** | 0.115 | **0.293** | **-0.380** | **-0.268** | **0.120** |
| Centriole position | 0.115 | **1** | -0.017 | 0.019 | -0.087 | -0.018 |
| Cell area | **0.293** | -0.017 | **1** | **-0.478** | **-0.390** | **0.629** |
| Cell height | **-0.380** | 0.019 | **-0.478** | **1** | **0.714** | **-0.344** |
| Cell circularity | **-0.268** | -0.087 | **-0.390** | **0.714** | **1** | **-0.271** |
| Nucleus area | **0.120** | -0.018 | **0.629** | **-0.344** | **-0.271** | **1** |

*P*-values (Pearson):

| Variables | Cilia length | Centriole position | Cell area | Cell height | Cell circularity | Nucleus area |
|---|---|---|---|---|---|---|
| Cilia length | **0** | 0.333 | **0.012** | **0.001** | **0.022** | **0.002** |
| Centriole position | 0.333 | **0** | 0.887 | 0.848 | 0.465 | 0.880 |
| Cell area | **0.012** | 0.887 | **0** | **<0.001** | **0.001** | **<0.001** |
| Cell height | **0.001** | 0.848 | **<0.001** | **0** | **<0.001** | **0.003** |
| Cell circularity | **0.022** | 0.465 | **0.001** | **<0.001** | **0** | **0.020** |
| Nucleus area | **0.002** | 0.880 | **<0.001** | **0.003** | **0.020** | **0** |

Coefficients of determination (Pearson):

| Variables | Cilia length | Centriole position | Cell area | Cell height | Cell circularity | Nucleus area |
|---|---|---|---|---|---|---|
| Cilia length | **1** | 0.013 | **0.086** | **0.144** | **0.048** | **0.130** |
| Centriole position | 0.013 | **1** | 0.000 | 0.000 | 0.008 | 0.000 |
| Cell area | **0.086** | 0.000 | **1** | **0.228** | **0.152** | **0.395** |
| Cell height | **0.144** | 0.000 | **0.228** | **1** | **0.509** | **0.118** |
| Cell circularity | **0.048** | 0.008 | **0.152** | **0.509** | **1** | **0.073** |
| Nucleus area | **0.130** | 0.000 | **0.395** | **0.118** | **0.073** | **1** |

*Shaded values in bold are different from 0 with a significance level alpha=0.05*

**Table S5.** Correlation test summary of 24 hours, 50 kPa

Summary statistics (Quantitative data):

| Variable | Observations | Minimum | Maximum | Mean | Std. deviation |
|---|---|---|---|---|---|
| Cilia length | 69 | 1.180 | 4.860 | 2.794 | 0.745 |
| Centriole position | 69 | 0.000 | 100.000 | 55.324 | 33.656 |
| Cell area | 69 | 193.750 | 3187.163 | 766.467 | 585.294 |
| Cell height | 69 | 1.406 | 4.580 | 2.486 | 0.663 |
| Cell circularity | 69 | 0.022 | 0.283 | 0.096 | 0.047 |
| Nucleus area | 69 | 47.563 | 713.190 | 208.795 | 116.711 |

Correlation matrix (Pearson):

| Variables | Cilia length | Centriole position | Cell area | Cell height | Cell circularity | Nucleus area |
|---|---|---|---|---|---|---|
| Cilia length | **1** | 0.128 | **0.319** | -0.155 | 0.008 | 0.189 |
| Centriole position | 0.128 | **1** | 0.047 | 0.005 | 0.106 | 0.052 |
| Cell area | **0.319** | 0.047 | **1** | **-0.260** | **-0.242** | **0.625** |
| Cell height | -0.155 | 0.005 | **-0.260** | **1** | 0.048 | -0.014 |
| Cell circularity | 0.008 | 0.106 | **-0.242** | 0.048 | **1** | **-0.283** |
| Nucleus area | 0.189 | 0.052 | **0.625** | -0.014 | **-0.283** | **1** |

*P*-values (Pearson):

| Variables | Cilia length | Centriole position | Cell area | Cell height | Cell circularity | Nucleus area |
|---|---|---|---|---|---|---|
| Cilia length | **0** | 0.294 | **0.008** | 0.202 | 0.945 | 0.119 |
| Centriole position | 0.294 | **0** | 0.700 | 0.967 | 0.388 | 0.671 |
| Cell area | **0.008** | 0.700 | **0** | **0.031** | **0.045** | **<0.001** |
| Cell height | 0.202 | 0.967 | **0.031** | **0** | 0.555 | 0.907 |
| Cell circularity | 0.945 | 0.388 | **0.045** | 0.555 | **0** | **0.019** |
| Nucleus area | 0.119 | 0.671 | **<0.001** | 0.907 | **0.019** | **0** |

Coefficients of determination (Pearson):

| Variables | Cilia length | Centriole position | Cell area | Cell height | Cell circularity | Nucleus area |
|---|---|---|---|---|---|---|
| Cilia length | **1** | 0.016 | **0.102** | 0.024 | 0.000 | 0.012 |
| Centriole position | 0.016 | **1** | 0.002 | 0.000 | 0.011 | 0.003 |
| Cell area | **0.102** | 0.002 | **1** | **0.068** | **0.059** | **0.390** |
| Cell height | 0.024 | 0.000 | **0.068** | **1** | 0.005 | 0.000 |
| Cell circularity | 0.000 | 0.011 | **0.059** | 0.005 | **1** | **0.080** |
| Nucleus area | 0.012 | 0.003 | **0.390** | 0.000 | **0.080** | **1** |

*Shaded values in bold are different from 0 with a significance level alpha=0.05*

**Table S6.** Correlation test summary of 48 hours, 50 kPa

Summary statistics (Quantitative data):

| Variable | Observations | Minimum | Maximum | Mean | Std. deviation |
|---|---|---|---|---|---|
| Cilia length | 66 | 0.862 | 3.470 | 2.222 | 0.644 |
| Centriole position | 66 | 0.000 | 100.000 | 51.758 | 32.874 |
| Cell area | 66 | 98.010 | 1625.770 | 506.978 | 333.198 |
| Cell height | 66 | 1.140 | 6.180 | 2.848 | 1.022 |
| Cell circularity | 66 | 0.010 | 0.940 | 0.199 | 0.244 |
| Nucleus area | 66 | 38.770 | 841.200 | 182.080 | 141.071 |

Correlation matrix (Pearson):

| Variables | Cilia length | Centriole position | Cell area | Cell height | Cell circularity | Nucleus area |
|---|---|---|---|---|---|---|
| Cilia length | **1** | 0.095 | 0.212 | -0.171 | -0.184 | 0.222 |
| Centriole position | 0.095 | **1** | -0.086 | 0.052 | -0.081 | 0.144 |
| Cell area | 0.212 | -0.086 | **1** | -0.101 | **-0.455** | **0.737** |
| Cell height | -0.171 | 0.052 | -0.101 | **1** | **0.323** | -0.101 |
| Cell circularity | -0.184 | -0.081 | **-0.455** | **0.323** | **1** | **-0.346** |
| Nucleus area | 0.222 | 0.144 | **0.737** | -0.101 | **-0.346** | **1** |

*P*-values (Pearson):

| Variables | Cilia length | Centriole position | Cell area | Cell height | Cell circularity | Nucleus area |
|---|---|---|---|---|---|---|
| Cilia length | **0** | 0.424 | 0.088 | 0.169 | 0.138 | 0.073 |
| Centriole position | 0.424 | **0** | 0.494 | 0.678 | 0.516 | 0.249 |
| Cell area | 0.088 | 0.494 | **0** | 0.419 | **<0.001** | **<0.001** |
| Cell height | 0.169 | 0.678 | 0.419 | **0** | **0.008** | 0.420 |
| Cell circularity | 0.138 | 0.516 | **<0.001** | **0.008** | **0** | **0.004** |
| Nucleus area | 0.073 | 0.249 | **<0.001** | 0.420 | **0.004** | **0** |

Coefficients of determination (Pearson):

| Variables | Cilia length | Centriole position | Cell area | Cell height | Cell circularity | Nucleus area |
|---|---|---|---|---|---|---|
| Cilia length | **1** | 0.009 | 0.045 | 0.029 | 0.034 | 0.049 |
| Centriole position | 0.009 | **1** | 0.007 | 0.003 | 0.007 | 0.021 |
| Cell area | 0.045 | 0.007 | **1** | 0.010 | **0.207** | **0.543** |
| Cell height | 0.029 | 0.003 | 0.010 | **1** | **0.104** | 0.010 |
| Cell circularity | 0.034 | 0.007 | **0.207** | **0.104** | **1** | **0.120** |
| Nucleus area | 0.049 | 0.021 | **0.543** | 0.010 | **0.120** | **1** |

*Shaded values in bold are different from 0 with a significance level alpha=0.05*

**Table S7.** Correlation test summary of 12 hours, coverslips (GPa)

Summary statistics (Quantitative data):

| Variable | Observations | Minimum | Maximum | Mean | Std. deviation |
|---|---|---|---|---|---|
| Cilia length | 85 | 0.590 | 5.000 | 2.579 | 0.810 |
| Centriole position | 85 | 0.000 | 78.000 | 17.565 | 26.124 |
| Cell area | 85 | 141.120 | 2400.270 | 775.747 | 452.617 |
| Cell height | 85 | 0.830 | 7.480 | 2.909 | 1.434 |
| Cell circularity | 85 | 0.030 | 0.710 | 0.162 | 0.140 |
| Nucleus area | 85 | 63.430 | 791.510 | 211.484 | 133.924 |

Correlation matrix (Pearson):

| Variables | Cilia length | Centriole position | Cell area | Cell height | Cell circularity | Nucleus area |
|---|---|---|---|---|---|---|
| Cilia length | **1** | **-0.383** | **0.302** | 0.062 | -0.074 | **0.346** |
| Centriole position | **-0.383** | **1** | -0.054 | **0.287** | 0.135 | **-0.217** |
| Cell area | **0.302** | -0.054 | **1** | 0.071 | -0.017 | **0.656** |
| Cell height | 0.062 | **0.287** | 0.071 | **1** | 0.106 | **-0.240** |
| Cell circularity | -0.074 | 0.135 | -0.017 | 0.106 | **1** | **-0.289** |
| Nucleus area | **0.346** | **-0.217** | **0.656** | **-0.240** | **-0.289** | **1** |

*P*-values (Pearson):

| Variables | Cilia length | Centriole position | Cell area | Cell height | Cell circularity | Nucleus area |
|---|---|---|---|---|---|---|
| Cilia length | **0** | **<0.001** | **0.005** | 0.570 | 0.501 | **<0.001** |
| Centriole position | **<0.001** | **0** | 0.624 | **0.008** | 0.220 | **0.046** |
| Cell area | **0.005** | 0.624 | **0** | 0.518 | 0.878 | **<0.001** |
| Cell height | 0.570 | **0.008** | 0.518 | **0** | 0.332 | **0.027** |
| Cell circularity | 0.501 | 0.220 | 0.878 | 0.332 | **0** | **0.007** |
| Nucleus area | **<0.001** | **0.046** | **<0.001** | **0.027** | **0.007** | **0** |

Coefficients of determination (Pearson):

| Variables | Cilia length | Centriole position | Cell area | Cell height | Cell circularity | Nucleus area |
|---|---|---|---|---|---|---|
| Cilia length | **1** | **0.147** | **0.091** | 0.004 | 0.005 | **0.120** |
| Centriole position | **0.147** | **1** | 0.003 | **0.083** | 0.018 | **0.047** |
| Cell area | **0.091** | 0.003 | **1** | 0.005 | 0.000 | **0.430** |
| Cell height | 0.004 | **0.083** | 0.005 | **1** | 0.011 | **0.058** |
| Cell circularity | 0.005 | 0.018 | 0.000 | 0.011 | **1** | **0.083** |
| Nucleus area | **0.120** | **0.047** | **0.430** | **0.058** | **0.083** | **1** |

*Shaded values in bold are different from 0 with a significance level alpha=0.05*

**Table S8.** Correlation test summary of 24 hours, coverslips (GPa)

Summary statistics (Quantitative data):

| Variable | Observations | Minimum | Maximum | Mean | Std. deviation |
|---|---|---|---|---|---|
| Cilia length | 79 | 1.640 | 8.650 | 3.755 | 1.745 |
| Centriole position | 79 | 0.000 | 77.000 | 12.430 | 21.879 |
| Cell area | 79 | 292.830 | 5382.100 | 1624.154 | 1198.138 |
| Cell height | 79 | 2.120 | 8.890 | 4.123 | 1.416 |
| Cell circularity | 79 | 0.030 | 0.510 | 0.113 | 0.096 |
| Nucleus area | 79 | 95.090 | 1859.880 | 494.909 | 437.966 |

Correlation matrix (Pearson):

| Variables | Cilia length | Centriole position | Cell area | Cell height | Cell circularity | Nucleus area |
|---|---|---|---|---|---|---|
| Cilia length | **1** | -0.002 | **0.591** | 0.123 | **-0.349** | **0.698** |
| Centriole position | -0.002 | **1** | 0.213 | 0.032 | -0.024 | 0.181 |
| Cell area | **0.591** | 0.213 | **1** | -0.124 | -0.216 | **0.903** |
| Cell height | 0.123 | 0.032 | -0.124 | **1** | 0.073 | 0.063 |
| Cell circularity | **-0.349** | -0.024 | -0.216 | 0.073 | **1** | **-0.311** |
| Nucleus area | **0.698** | 0.181 | **0.903** | 0.063 | **-0.311** | **1** |

*P*-values (Pearson):

| Variables | Cilia length | Centriole position | Cell area | Cell height | Cell circularity | Nucleus area |
|---|---|---|---|---|---|---|
| Cilia length | **0** | 0.983 | **<0.001** | 0.282 | **0.002** | **<0.001** |
| Centriole position | 0.983 | **0** | 0.060 | 0.779 | 0.673 | 0.111 |
| Cell area | **<0.001** | 0.060 | **0** | 0.275 | 0.056 | **<0.001** |
| Cell height | 0.282 | 0.779 | 0.275 | **0** | 0.521 | 0.581 |
| Cell circularity | **0.002** | 0.673 | 0.056 | 0.521 | **0** | **0.005** |
| Nucleus area | **<0.001** | 0.111 | **<0.001** | 0.581 | **0.005** | **0** |

Coefficients of determination (Pearson):

| Variables | Cilia length | Centriole position | Cell area | Cell height | Cell circularity | Nucleus area |
|---|---|---|---|---|---|---|
| Cilia length | **1** | 0.000 | **0.350** | 0.015 | **0.122** | **0.247** |
| Centriole position | 0.000 | **1** | 0.045 | 0.001 | 0.002 | 0.033 |
| Cell area | **0.350** | 0.045 | **1** | 0.015 | 0.047 | **0.815** |
| Cell height | 0.015 | 0.001 | 0.015 | **1** | 0.005 | 0.004 |
| Cell circularity | **0.122** | 0.002 | 0.047 | 0.005 | **1** | **0.097** |
| Nucleus area | **0.247** | 0.033 | **0.815** | 0.004 | **0.097** | **1** |

*Shaded values in bold are different from 0 with a significance level alpha=0.05*

**Table S9.** Correlation test summary of 48 hours, coverslips (GPa)

Summary statistics (Quantitative data):

| Variable | Observations | Minimum | Maximum | Mean | Std. deviation |
|---|---|---|---|---|---|
| Cilia length | 90 | 0.870 | 4.780 | 2.658 | 0.774 |
| Centriole position | 90 | 0.000 | 100.000 | 14.578 | 23.571 |
| Cell area | 90 | 141.120 | 2526.080 | 715.698 | 442.896 |
| Cell height | 90 | 1.110 | 6.470 | 2.968 | 1.374 |
| Cell circularity | 90 | 0.030 | 0.650 | 0.195 | 0.158 |
| Nucleus area | 90 | 66.510 | 762.590 | 223.412 | 128.884 |

Correlation matrix (Pearson):

| Variables | Cilia length | Centriole position | Cell area | Cell height | Cell circularity | Nucleus area |
|---|---|---|---|---|---|---|
| Cilia length | **1** | **-0.214** | **0.334** | 0.165 | -0.043 | **0.265** |
| Centriole position | **-0.214** | **1** | -0.153 | -0.133 | **-0.221** | -0.026 |
| Cell area | **0.334** | -0.153 | **1** | -0.003 | 0.053 | **0.486** |
| Cell height | 0.165 | -0.133 | -0.003 | **1** | 0.092 | **-0.246** |
| Cell circularity | -0.043 | **-0.221** | 0.053 | 0.092 | **1** | **-0.292** |
| Nucleus area | **0.265** | -0.026 | **0.486** | **-0.246** | **-0.292** | **1** |

*P*-values (Pearson):

| Variables | Cilia length | Centriole position | Cell area | Cell height | Cell circularity | Nucleus area |
|---|---|---|---|---|---|---|
| Cilia length | **0** | **0.042** | **0.001** | 0.121 | 0.691 | **0.012** |
| Centriole position | **0.042** | **0** | 0.149 | 0.213 | **0.012** | 0.805 |
| Cell area | **0.001** | 0.149 | **0** | 0.979 | 0.622 | **<0.001** |
| Cell height | 0.121 | 0.213 | 0.979 | **0** | 0.387 | **0.019** |
| Cell circularity | 0.691 | **0.012** | 0.622 | 0.387 | **0** | **0.005** |
| Nucleus area | **0.012** | 0.805 | **<0.001** | **0.019** | **0.005** | **0** |

Coefficients of determination (Pearson):

| Variables | Cilia length | Centriole position | Cell area | Cell height | Cell circularity | Nucleus area |
|---|---|---|---|---|---|---|
| Cilia length | **1** | **0.046** | **0.112** | 0.027 | 0.002 | **0.070** |
| Centriole position | **0.046** | **1** | 0.023 | 0.018 | **0.049** | 0.001 |
| Cell area | **0.112** | 0.023 | **1** | 0.000 | 0.003 | **0.526** |
| Cell height | 0.027 | 0.018 | 0.000 | **1** | 0.009 | **0.060** |
| Cell circularity | 0.002 | **0.049** | 0.003 | 0.009 | **1** | **0.086** |
| Nucleus area | **0.070** | 0.001 | **0.526** | **0.060** | **0.086** | **1** |

*Shaded values in bold are different from 0 with a significance level alpha=0.05*
